# Harnessing landscape genomics to evaluate genomic vulnerability and future climate resilience in an East Asia perennial

**DOI:** 10.1101/2025.01.19.633819

**Authors:** Li Feng, Cong-Ying Wang, Li-Pan Zhou, Yi-Han Wang, Jing Wang, Zheng-Yuan Wang, Tao Zhou, Xu-Mei Wang

## Abstract

In this era of rapid climate change, understanding the adaptive potential of organisms is imperative for buffering biodiversity loss. Genomic forecasting provides invaluable insights into population vulnerability and adaptive potential under diverse climatic conditions, thereby facilitating management interventions, bolstering population resilience, and shaping germplasm conservation strategies tailored to specific species. Here we integrated population genomics and landscape genomics approaches, leveraging single-nucleotide polymorphisms obtained through whole-genome resequencing of 43 *Rheum palmatum* complex populations, to pinpoint adaptive variation and its significance in the context of future climates, delineate seed zones, and establish guidelines for *ex situ* germplasm conservation to capture the majority of existing adaptive diversity. Our analysis unveiled that the species complex comprised two distinct genetic clusters, exhibiting differential climate adaptation and genomic vulnerabilities across its distribution range. We also determined that the species range could be subdivided into three distinct seed zones, with varying sample requirements per seed zone corresponding to differed conservation efforts. Overall, our findings provide a genome-wide perspective on climate adaptation and valuable insights into germplasm conservation strategies aimed at enhancing population resilience in future climates, serving as a blueprint for restoration plans of other vulnerable species.

## Introduction

In this era of rapid climate change, a profound understanding of organisms’ adaptive potential is paramount for advancing proactive conservation efforts and mitigating biodiversity loss (Hoffmann and Sgrò 2011; Hoffmann, et al. 2021). Numerous studies have indicated that many species are unlikely to adapt to climatic change by shifting their geographical distribution; instead, their adaptation relies on phenotypic plasticity and evolutionary adaptation (Scheffers, et al. 2016; Aitken, et al. 2024). As a result, climate adaptation strategies have become crucial for preserving and managing wild populations, to ensure an appropriate and swift response to the climatic alterations (Forester, et al. 2022). These strategies must prioritize the identification or augmentation of species’ or populations’ evolutionary potential and adaptive capabilities, grounded in established or hypothesized pathway(s) and mechanism(s) through which climate impacts the focal taxa (Bay, et al. 2018; Exposito-Alonso, et al. 2019). Among the diverse methodologies available, genomic forecasting has emerged as a particularly cost-effective technique for predicting future disruptions to locally adaptive gene-environment associations (Capblancq, Fitzpatrick, et al. 2020). This approach offers invaluable insights into population vulnerability and adaptive potential across various environments, enabling informed targeted management interventions, the enhancement of population resilience, and the development of species-specific germplasm conservation strategies (Capblancq, Fitzpatrick, et al. 2020; Bernatchez, et al. 2024).

Genomic forecasting, which commonly employs genomic offset (alternatively termed genetic offset (Fitzpatrick and Keller 2015) or genomic vulnerability (Bay, et al. 2018)), has found applications in various taxa (Gougherty, et al. 2021; Layton, et al. 2021; Chen, et al. 2023). This method involves mapping the current spatial distribution of adaptive alleles onto existing environmental conditions, and subsequently computing the disparity between the present and future climate-associated genomic composition as an indicator of climate change maladaptation (Fitzpatrick and Keller 2015). Nevertheless, solely relying on genomic forecasting is insufficient for guiding adaptive management interventions and enhancing the resilience of species possessing significant ecological and economic value. Identifying the genomic targets of climate-related selection stands as a pivotal step in comprehending local adaptation within these species, and in formulating strategies for preserving pertinent genomic diversity. One approach to achieving this aim is through the lens of seed zones, where a species range is partitioned into the minimal number of regions that effectively delineate climate and ecoregions (Sandercock, et al. 2024).

Seed zones, which accommodate local adaptation and population variation, are extensively employed for germplasm management and conservation, with the primary goal of preventing the utilization of maladaptive seed sources in order to bolster population resilience in future (Hufford and Mazer 2003; Yu, et al. 2022). Traditionally, seed zones have partitioned the entire species range into distinct regions, with the allocation of seeds confined within each designated zone, following the “local is best” paradigm (Broadhurst, et al. 2008). This presumption is based on the belief that existing populations are well-acclimated to their local conditions. Nevertheless, as climate change progresses, the concept of a static “local climate” is becoming challenged, leading to a mismatch between the climatic conditions that local populations have historically adapted to and the future climatic scenarios they are destined to encounter (Aitken, et al. 2008). Climate change has induced swift alterations in preexisting habitat conditions, thus presenting novel challenges in aligning appropriate planting environments with suitable seed sources (Harrison 2021). Consequently, the development of climate-driven seed transfer models becomes imperative, facilitating the implementation of genetically informed management strategies, such as assisted gene flow (Aitken and Whitlock 2013). Despite the implementation of climate-based seed transfer practices, there remains a necessity to classify populations according to their comparable adaptive diversity, thereby facilitating streamlined management and deployment procedures. As genomic and environmental data rapidly accumulate and become easily accessible, landscape genomic methods are arising as a viable alternative for geographically delineating seed zones, vital for managing adaptive variation and breeding (Yu, et al. 2022; Sandercock, et al. 2024). Based on this information provided by landscape genomics, we are able to assess the necessary sampling intensity to capture the existing adaptive diversity within the designated seed zones, constituting a logical step towards enhancing population resilience in the face of future climate changes.

The *Rheum palmatum* complex, comprising *R. officinale*, *R. palmatum*, and *R. tanguticum*, predominantly occurs at mid- to high-altitudes (*ca*. 1,000-5,100 m above sea level) within the mountainous areas of western China (Bao and Grabovskaya-Borodina 2003). These regions are distinguished by their exceptional biodiversity and endemism but are also highly vulnerable to the impacts of climate change (Myers, et al. 2000; Rahbek, et al. 2019). Rhubarb derived from the dried roots and rhizomes of these species and revered as the “king of herbs”, constitutes an important Traditional Chinese Medicine (TCM) that has been utilized for over 2,000 years as a purgative (Xiang, et al. 2020). Previous studies have uncovered that these herbs are likely to constitute a single species, with intraspecific lineages exhibiting a phylogeographical west-east split near/across the Sichuan Basin (Wang, et al. 2018; Feng, et al. 2020). These lineages often display confinement to ecologically and topologically heterogeneous habitats, thus suggesting potential signals of climate-driven selection. Therefore, this species complex provides an exemplary system for determining the intraspecific variability in genotype-climate associations and its resultant implications for climate change-induced vulnerability. To date, the majority of genomic vulnerability research has concentrated on trees (Gougherty, et al. 2021; Sang, et al. 2022; Sandercock, et al. 2024) and vertebrates (Bay, et al. 2018; Brauer, et al. 2023; Chen, et al. 2023). Although a growing number of studies have delved into the genomic vulnerability of herbs (Exposito-Alonso, et al. 2019; Zhao, et al. 2023; Zhang, Long, et al. 2024), a substantial knowledge deficit remains concerning the response of herbs to projected climate changes, especially given their occupancy of up to 40% of terrestrial areas and the considerable variability in their response to climatic stressors (Craine, et al. 2013; Petermann and Buzhdygan 2021).

Here, we describe the genomic basis of local adaptation within the species complex and present sampling recommendations aimed at bolstering germplasm conservation initiatives. By harnessing whole-genome resequencing data from 201 accessions of the species complex, we seek to: i) pinpoint genetic loci that exhibit significant associations with climatic variables; ii) quantitatively assess and geographically map the vulnerable populations in the context of future climatic shifts; iii) synthesize how these climate-linked loci collectively vary across landscapes, thereby enabling the delineation of seed zones critical for germplasm preservation; and iv) formulate sampling guidelines targeted at capturing adaptive diversity in wild populations. Our findings provide a comprehensive, genome-wide lens on climate adaptation within the *R. palmatum* complex, offering valuable insights into the progress of *ex situ* germplasm conservation efforts aimed at enhancing population resilience in the face of future climates.

## Results

### Genetic differentiation and population demography

Our comprehensive whole-genome resequencing effort has yielded approximately 14.30 million single nucleotide polymorphisms (SNPs) that successfully passed the initial quality filtering, derived from 213 individuals spanning the natural range of the species complex (Fig. 1; supplementary table S1). Notably, an average of approximately 94.7% of the clean reads were successfully aligned to the reference genome (2.8 Gb) of *R. palmatum* (Zhang, et al. 2024), achieving a mean depth of 18.9× and a coverage of 94.3% (supplementary table S2). After applying rigorous and stringent thresholds for SNP calling to ensure quality control and enable precise evaluation in subsequent statistical analyses (Marees, et al. 2018), we identified approximately 1.43 million high-quality SNPs from 201 samples of our focal species. Our findings revealed a high genetic differentiation (*F*_ST_) across all populations, with an average *F*_ST_ value of 0.22 (supplementary table S3). Furthermore, we observed variable levels of genetic diversity among these populations, exhibiting a mean observed heterozygosity (*H*_O_) of 0.19 and a mean heterozygosity (*H*_S_) of 0.13 within populations (supplementary table S1). Bayesian clustering utilizing 378,620 SNPs extracted from an LD-pruned dataset comprising 1.43 million SNPs identified a distinct separation between western and eastern populations emerged, with an optimal k-value of two (Fig. 1a, b; supplementary fig. S1a, b). Furthermore, the western cluster, primarily occupying the Hengduan mountains and adjacent areas, could be further subdivided into two distinct lineages (Fig. 1b; supplementary fig. S1c, d). It is worth mentioning that some individuals, particularly those belonging to western sub-lineages, exhibit evident admixture between sub-lineages. Both principal component analysis (PCA) and discriminant analysis of principal components (DAPC) consistently supported these findings (supplementary fig. S1e, f). Furthermore, our cluster analyses, which were based on pan-adaptation or core adaptation loci, yielded results that were comparable to those obtained through the utilization of 378,620 SNPs (supplementary fig. S2). Additionally, based on putatively neutral SNPs, we detected significant signals of isolation-by-distance (IBD) within the species complex (Fig. 1c). Collectively, these diverse lines of evidence strongly suggest that our focal species is likely to have originated from distinct refugia (Feng, et al. 2020).

**Fig. 1.**
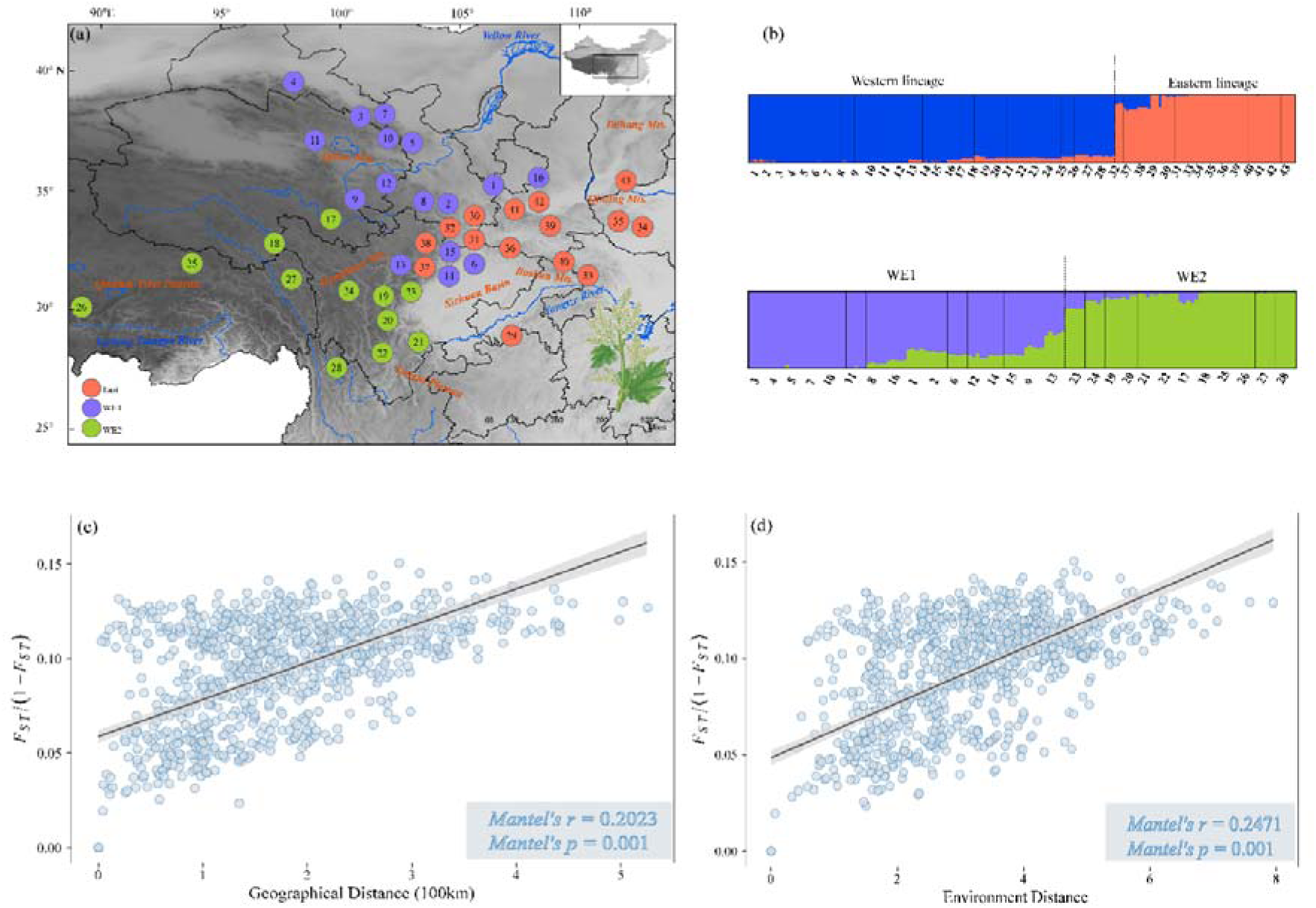
Population structure and genetic differentiation within the *Rheum palmatum* complex. (a) Geographical distribution of the 43 populations, with color-coded circles representing specific lineages identified by STRUCTURE. (b) Population assignment of 201 individuals based on STRUCTURE. Panels (c) and (d) represent the isolation-by-distance analysis (two-sided Mantel test) and isolation-by-environment analysis (two-sided partial Mantel test) for putative neutral loci, respectively, with shadows indicating the 95% confidence interval.

Subsequently, we utilized both the stairway plot and SMC++ to independently assess the changes in effective population size (*Ne*) for each of the three genetic lineages throughout their respective evolutionary trajectories (supplementary fig. S3a, b). Generally, the *Ne* trajectories inferred from the both methods provided evidence of a population decline across the three clusters, which was further substantiated by Tajima’s *D* statistics indicating positive *D* values for all populations (supplementary table S1). Ultimately, an analysis of linkage disequilibrium (LD) decay across the genome, taking into account physical distance, unveiled comparable trends among the three clusters. Specifically, the *r²* value decreased to below 0.1 at mean physical distances of approximately 5 kb and 1 kb for the eastern and western populations, respectively (supplementary fig. S3c). Collectively, these lines of evidence point to a potential recent divergence among the lineages within the species complex.

### Genomic signatures of local adaptation

We integrated genomic and climatic datasets to detect signatures of divergent selection by employing genotype-environment association (GEA) methodologies. Initially, we utilized gradient forest (GF) (Ellis, et al. 2012), a machine-learning approach to pinpoint the climatic variables exhibiting the strongest association with genetic variation within the species complex. Among the 19 BIOCLIM variables (Fick and Hijmans 2017) tested (supplementary table S4), we selected the top six uncorrelated variables: mean diurnal range (BIO2), isothermality (BIO3), temperature seasonality (BIO4), mean temperature of the warmest quarter (BIO10), annual precipitation (BIO12), and precipitation seasonality (BIO15). These were retained as climate candidates based on their ranked importance and the correlations observed among the 19 climatic variables (supplementary fig. S4), ensuring the avoidance of multicollinearity issues in subsequent GEA analyses. To identify SNPs associated with the six climatic variables, we employed both the latent factor mixed model (LFMM) (Frichot, et al. 2013) and redundancy analysis (RDA) (Capblancq and Forester 2021). Through these analyses, we aimed to determine the genomic signatures underlying local adaptation in our focal species. Given that the overlapping detections of putatively adaptive SNPs across various methods may introduce biases towards the identification of loci undergoing strong selective sweeps and the RDA is capable of effectively detecting signatures of covarying multilocus selection (François, et al. 2016; Forester, et al. 2018), the comprehensive set of 16,709 putatively adaptive SNPs, identified through a combination of LFMM and RDA analyses, were collectively regarded as pan-adaptation loci (supplementary fig. S5a). Among the 10,988 significant adaptive SNPs identified via LFMM, 1,198 SNPs exhibited extreme loadings (with a standard deviation > 2.7) along one or multiple RDA axes, qualifying them as core adaptation loci. These two distinct categories of putatively adaptive loci are broadly distributed across the genome (supplementary fig. S5b). Functional enrichment analysis focusing on the genes associated with these loci revealed a significant enrichment in gene ontology (GO) categories related to biological processes and molecular function such as transcriptional regulation, DNA repair and response to cold (supplementary table S5), which are crucial for understanding the plant’s persistence under anticipated climate changes (Wellenreuther and Bernatchez 2018).

To further assess the validity of the SNPs identified in this study, we employed Mantel and partial Mantel tests to explore patterns of IBD and isolation-by-environment (IBE) for both potentially adaptive and neutral SNPs, respectively (Fig. 1c, d; supplementary fig. S6). Our findings revealed significant IBD patterns for both adaptive and neutral SNPs, albeit with stronger patterns observed for adaptive SNPs compared to neutral ones. Nonetheless, akin to IBD, both SNP types exhibited significant IBE patterns in partial Mantel tests, indicating that the genetic variation of adaptive SNPs was predominantly influenced by geography and environment. To disentangle the relative contributions of climate, geography, and population structure in explaining both adaptive and neutral genetic variation, we conducted partial RDA (pRDA) focused on pan-adaptation and core adaptation loci, respectively. The pan-adaptation loci analysis showed that, climate, geography, and neutral structure accounted for 78.3% of the variance in allele frequencies among localities of our focal species (supplementary table S6). Specifically, the exclusive contribution of climate effects was notable, explaining 9.8% of the total genetic variation (representing 12.5% of the variation explicable by the full model). The pure effect of neutral genetic structure was also significant, accounting for 16.2% of the total genetic variance (20.6% of the explicable variation), while geography contributed significantly to 2.3% (3.0% of the explicable variation). The results from pRDA centered on core adaptation loci mirrored those observation for pan-adaptation loci (supplementary tables S7). Interestingly, we discovered that a substantial portion of the explained genetic variance was entangled across various predictor groups, indicating a high level of collinearity among climate, geography, and neutral genetic structure.

### Spatial distribution of adaptive genomic diversity and genomic offset

Using the aggregate community-level turnover functions derived from putatively adaptive SNPs integrated within the GF framework and adaptively enriched RDA, we constructed a visual representation of the genotype-climate turnover surface spanning the potential ranges of the species complex. This visualization was achieved by utilizing the first two principal components extracted from the outputs of both the GF model and RDA. Our findings from the GF analysis indicated that the spatial turnover patterns were comparable between pan-adaptation and core adaptation loci, highlighting distinct genomic composition among populations within the species complex’s distribution range (Fig. 2). Furthermore, the adaptively enriched RDA conducted on the pan-adaptation and core adaptation loci delineated two principal gradients of climatic adaptation within our focal species (supplementary figs. S7-S8). The first axis of variation, denoted as RDA1, juxtaposed localities exhibiting a marked increase in isothermality (BIO3) – predominantly observed in the western regions of the species complex’s distribution range – against those experiencing greater temperature seasonality (BIO4), primarily situated in the central and eastern portions of its range. The second axis, RDA2, exhibited a stronger correlation with annual precipitation (BIO12) in low-elevation habitats, whereas it was associated with increased precipitation seasonality (BIO15) and a higher diurnal temperature range (BIO2) in western mountainous regions. These findings suggest distinct differences in genotype-climate associations between the western and eastern lineages, likely reflecting local adaptation to diverse climatic conditions.

**Fig. 2.**
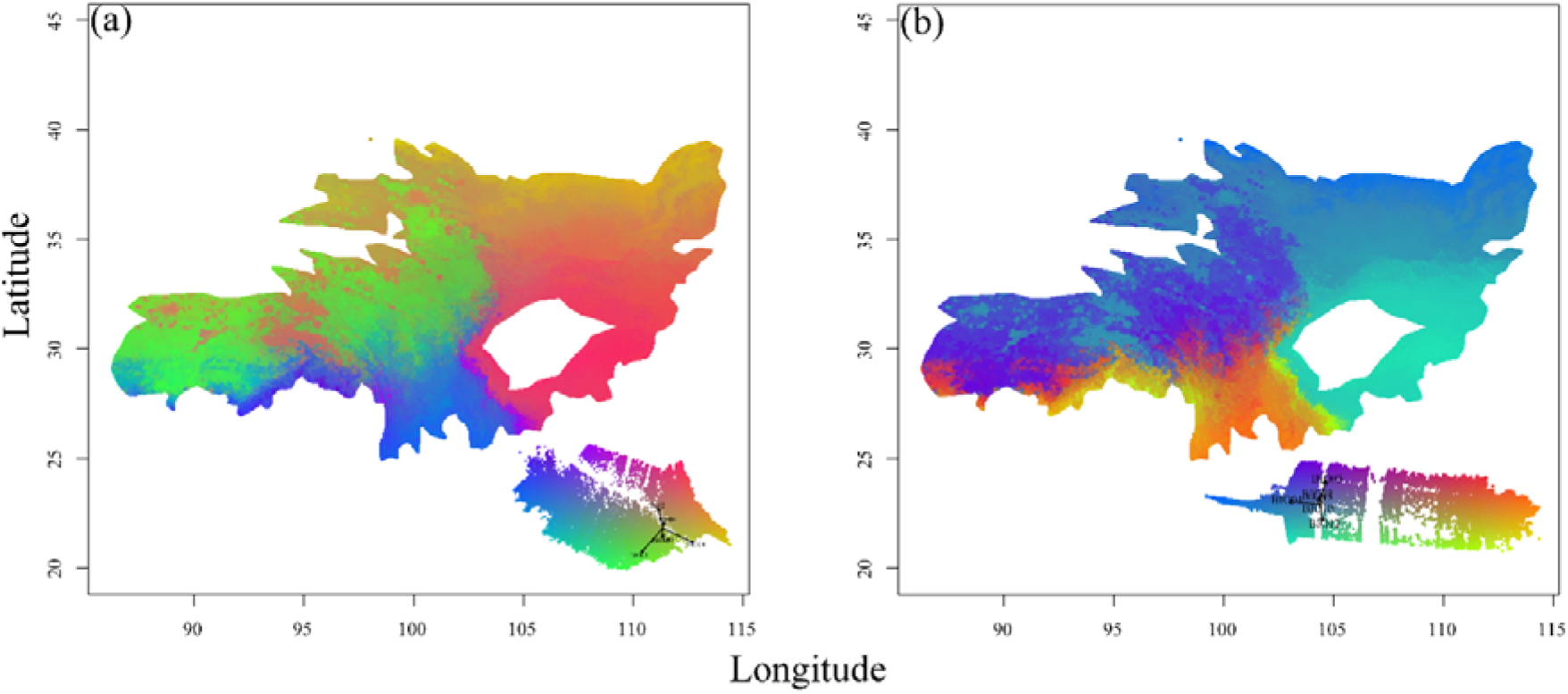
Predicted spatial variation in population-level genetic composition derived from gradient forest analysis for (a) pan-adaptation loci and (b) core adaptation loci. Regions shaded in similar colors are anticipated to host populations exhibiting comparable genetic composition. The biplot in each subfigure illustrates the influence of climate variables on the predicted genetic turnover, with labeled vectors highlighting the impact of the six top climate variables.

To anticipate how the spatial distribution of adaptive genetic variation might influence the long-term persistence of populations, we extrapolated the GEAs from GF model and RDA to future climate scenarios to estimate genomic offset. Populations exhibiting larger offsets are predicted to be more susceptible to climatic shifts. To accommodate uncertainties inherent in climate modeling, we forecasted genomic offset for the species complex using integrated projections from four distinct future climate models (BCC-CSM2-MR, ACCESS-CM2, MIROC6, and MPI-ESM1-2-HR) under both the lowest and highest greenhouse gas emission scenarios (SSP126 and SSP585) for the years 2050 (2041-2060) and 2090 (2081-2100). Extrapolations of genomic offset for the two SSPs in years 2050 and 2090, conducted through GF and RDA, unveiled analogous spatial patterns (Fig. 3; supplementary fig. S9). These patterns highlighted elevated risks of maladaptation in the central-eastern and southeastern portions of the species’ range, with diminished risks in the western regions. Notably, the estimations of genomic offset for SSP585 in 2090 using core adaptation loci via GF and RDA suggested that populations residing in certain southern sections of the western regions might maladapt to future climate changes (supplementary fig. S9). Moreover, projections of genomic offset utilizing both pan-adaptation and core adaptation loci across diverse SSPs and timeframes consistently demonstrated that populations within the eastern lineage exhibited higher offset values than those within the western lineage, with the exception of genomic offset estimations using core adaptation loci via RDA for SSP585 in 2090 (Fig. 4; supplementary fig. S10). Additionally, we conducted lineage-level predictions of variable importance using two datasets of putatively adaptive loci based on GF (supplementary fig. S11), and then estimated maladaptation regions, considering the potential for unrealistic estimations due to substantial genetic structure (Aguirre-Liguori, et al. 2023; Lind, et al. 2024). The genomic offset estimations based on GF at lineage levels uncovered consistent trends in estimating genomic offsets at the species level, and further revealed some fine-scale maladaptation zones arose within western regions under projected climate changes. (Figs. 3a-d, 5 and supplementary fig. S9a-d). However, genomic offset estimations at lineage levels using RDA were inconsistent with the results obtained at the species level (Fig. 3e-h, supplementary figs. S9e-h and S12). Further exploration is imperative to elucidate the impact of substantial genetic structure on the projection of maladaptation regions when employing RDA. Nonetheless, our results indicate that distinct GEAs between the western and eastern lineages within the distribution range of our focal species are potentially facilitating estimations of fine-scale maladaptation regions compared to those at the species level in response to impending climatic perturbations.

**Fig. 3.**
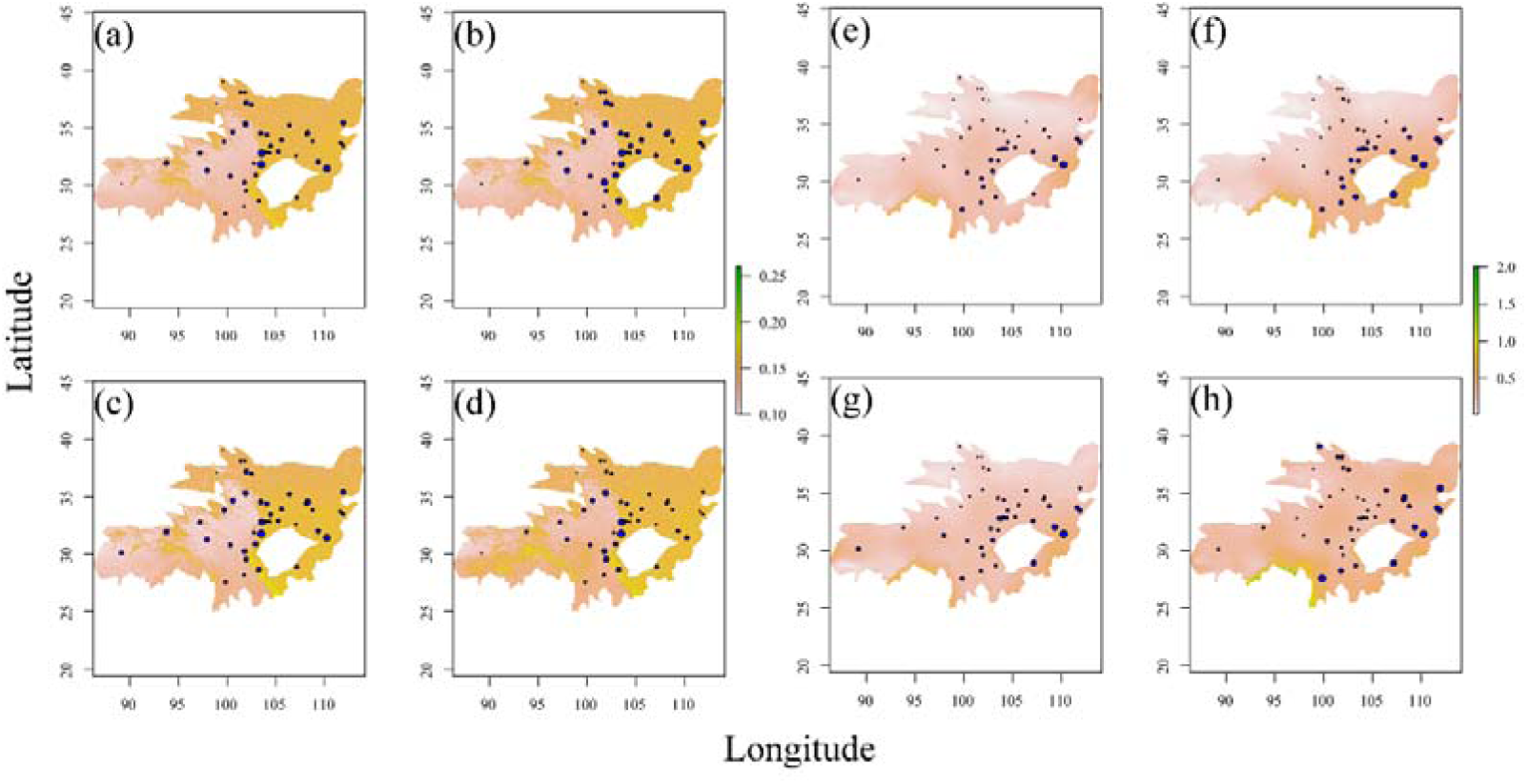
Predicted genomic offsets across the range of *Rheum palmatum* complex in response to future climates, derived from pan-adaptation loci using gradient forest (panels a-d) and redundancy analysis (panels e-h). Panels (a) and (e) represent the scenario of shared socioeconomic pathways SSP126 in 2050, while (b) and (f) depict SSP126 in 2090. Similarly, (c) and (g) illustrate SSP 585 in 2050, and (d) and (h) show SSP585 in 2090. The size of the blue circles in each panel corresponds to the magnitude of the genomic offset under the respective climate scenarios.

**Fig. 4.**
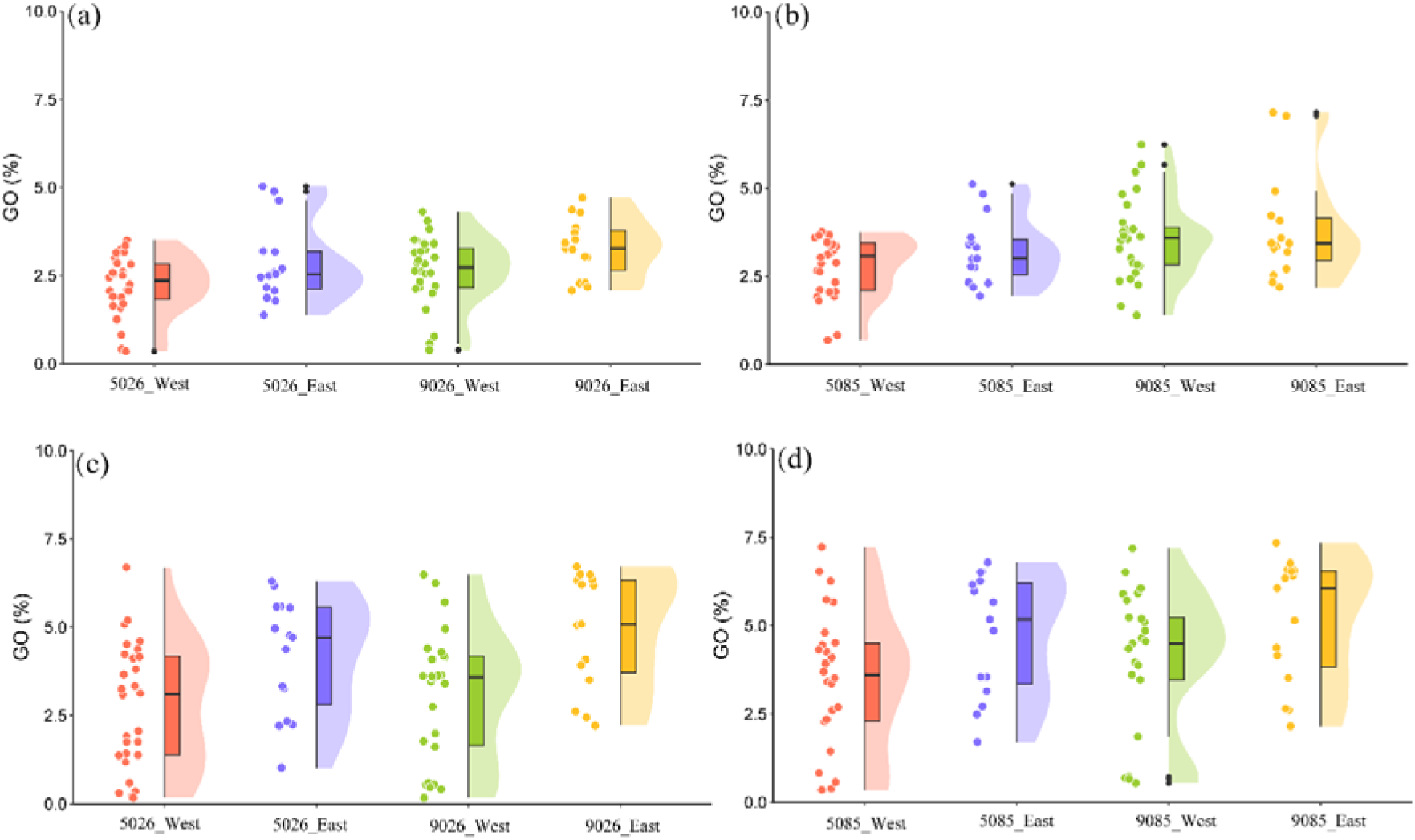
Predicted genomic offsets of the western and eastern lineage populations derived from species-level estimations, utilizing both pan-adaption loci (panels a and b) and core adaptation loci (panels c and d) via gradient forest. On the x-axis, the numerals 5026 and 9026 denote the scenarios corresponding to the shared socioeconomic pathways SSP126 in 2050 and 2090, respectively, while 5085 and 9085 represent the SSP585 scenarios in 2050 and 2090, respectively.

**Fig. 5.**
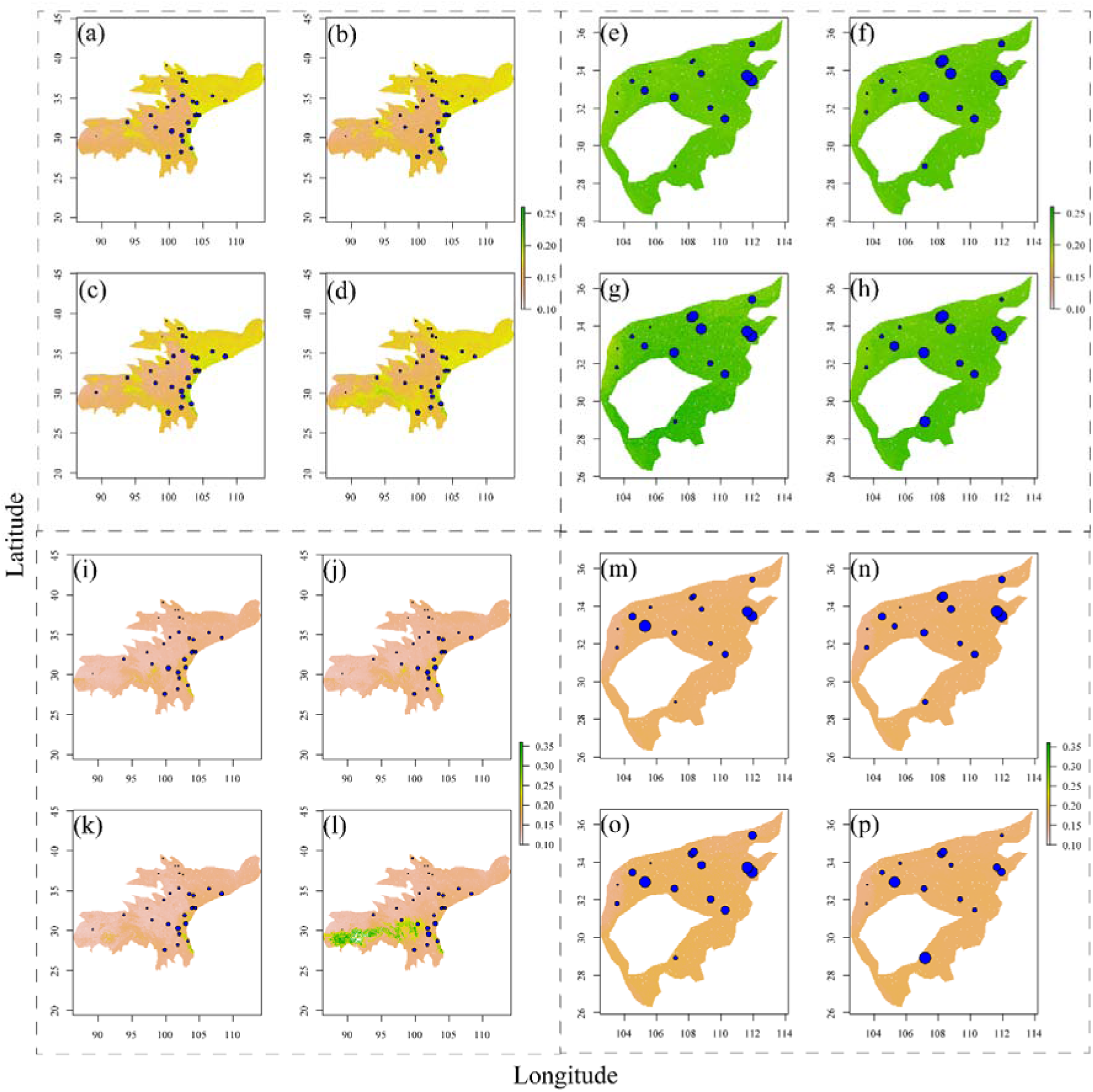
Predicted genomic offsets at the lineage level based on pan-adaptation loci (panels a-h) and core adaptation loci (panels i-p) using gradient forest. Panels (a) and (i) depict the offsets for western lineage populations under the scenarios corresponding to the shared socioeconomic pathways SSP126 in 2050, while (b) and (j) represent the same for 2090. Panels (c) and (k) exhibit offsets for SSP585 in 2050, and (d) and (l) for 2090. For eastern lineage populations, panels (e) and (m) display offsets under SSP126 in 2050, (f) and (n) for 2090, (g) and (o) for SSP585 in 2050, and finally, (h) and (p) for 2090.

We ultimately investigated the correlations between genomic offset, genetic diversity, and genetic load, aiming to ascertain whether populations exhibiting a higher genetic offset also bear an augmented burden of deleterious mutations or diminished genetic diversity. Typically, populations with elevated genetic load maybe more susceptible to climatic shifts compared to those with lower genetic load (Nadeau and Urban 2019; Aguirre-Liguori, et al. 2021). Utilizing SIFT4G (Vaser, et al. 2016), we anticipated and categorized coding SNPs into four distinct groups based on their impact: synonymous, tolerated, deleterious, and loss of function (LOF). To approximate genetic load, we employed the ratios of derived functional variants (comprising tolerated, deleterious, or LOF mutations) to synonymous variants. The results disclosed no discernible patterns of association between the anticipated genomic offsets and genetic loads, a finding that remained consistent even when considering LOF variants presumed to exert substantial deleterious effects (supplementary figs. S13-S16). Additionally, our results also elucidated that, in the majority of instances, populations exhibiting high genomic offsets typically possessed low genetic diversity, lending support to the conclusion that genetic diversity is proximate to adaptive potential and crucial for sustaining species’ adaptation (Scott, et al. 2020; Exposito-Alonso, et al. 2022). Whether this relationship holds true across other species remains to be rigorously evaluated in future studies within a meticulously designed framework.

### Seed zone delineation, range shift prediction and sampling recommendations

Given that climate served as a primary determinant of genomic variation within our focal species, we postulated the feasibility of delineating seed zones, whose boundaries would reflect precipitation and temperature gradients within the potential range of the species complex. The principal components, representing the continuous genomic variation predicted by GF, can be clustered into varying numbers of classes to delineate seed zones, thereby enabling *ex situ* germplasm conservation that facilitates climate-matched population resilience (Yu, et al. 2022; Sandercock, et al. 2024). We determined that three zones were optimal for both datasets of putatively adaptive loci (supplementary figs. S17). A longitudinal boundary situated at approximately 103°E and a latitudinal boundary positioned at roughly 28°N delineated the species’ range into three distinct seed zones: SZ1 and SZ3, both located in the western region, and SZ2, situated in the eastern region (Fig. 6a). SZ1 was heavily influenced by BIO03, whereas SZ2 exhibited a strong influence from BIO04 (Fig. 6b). To mitigate the effects of climate change, the geographic centroids of these seed zones required shifts ranging from approximately 48.3 km to 359.3 km from their current ranges, as estimated by genomic offsets derived from pan-adaptation loci under low and high climate change scenarios (Fig. 6c-f). Additionally, the SZ2 centroid was predicted to necessitate the greatest average shift among all scenarios, followed by SZ3 and SZ1. The outcomes of climate-driven seed zone shift prediction using core adaptation loci echoed those points for pan-adaptation loci (supplementary fig. S18).

**Fig. 6.**
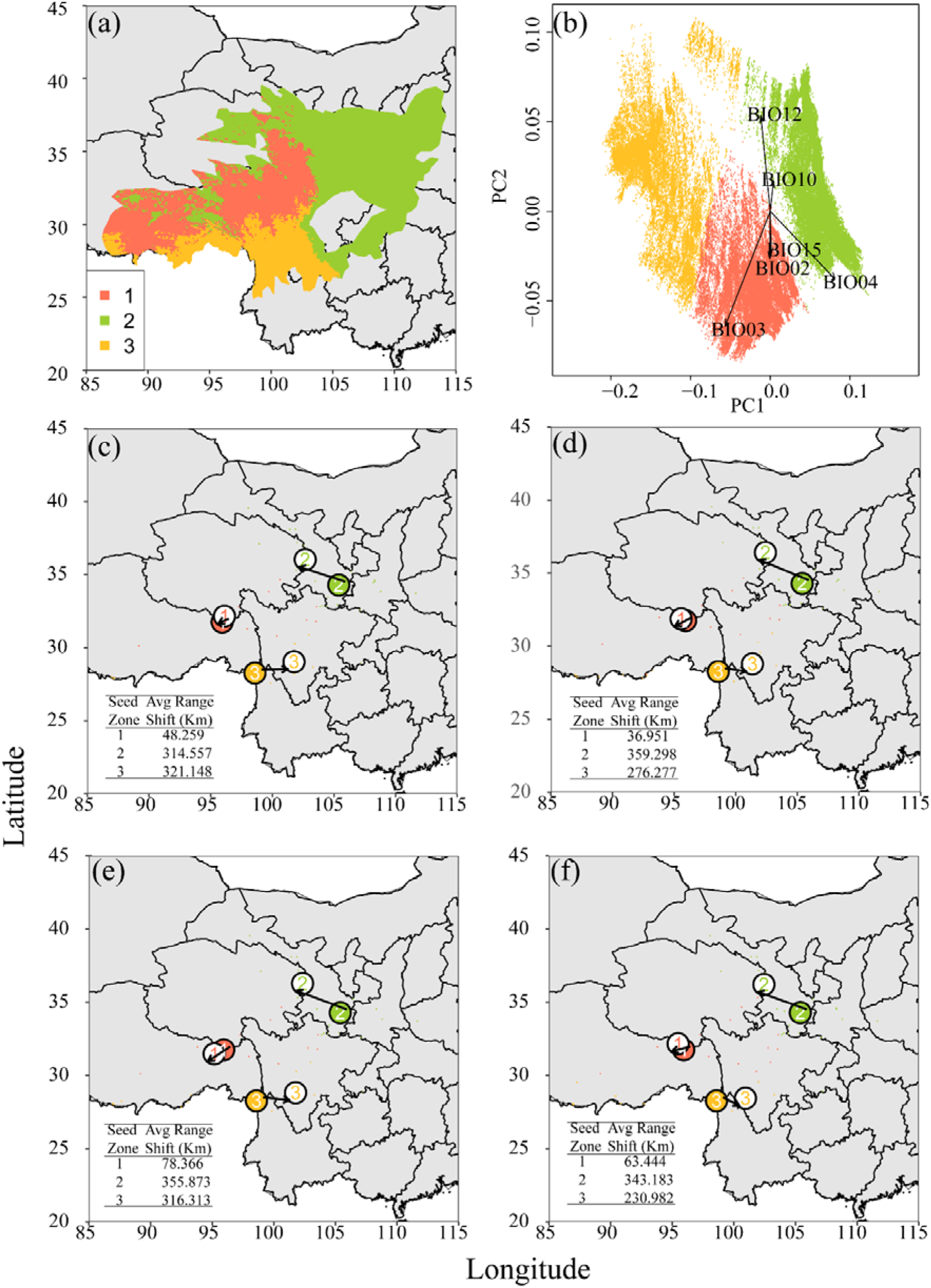
Predicted seed zones based on pan-adaptation loci and their range shifts to minimize genomic offset under climate change. Panel (a) represents predicted three seed zones for the *Rheum palmatum* complex using gradient forest (GF). The biplot in subfigure illustrates the GF-predicted genomic variation used for seed zone delineation. Panels (b-e) represent the predicted range adjustments for each seed zone to minimize genomic disparities derived from pan-adaptation loci under various climate scenarios. Solid-colored numerals denote the current centroids of the seed zones, whereas numerals on a white background indicate the anticipated centroid shifts in alignment with future climate projections. Panels (b) and (c) correspond to the SSP126 socioeconomic pathway scenarios in 2050 and 2090, respectively. Panels (d) and (e) reflect the SSP585 scenarios in 2050 and 2090, respectively. The colored pixels represent the predicted relocation points for each seed zone, where genomic disparities are minimized based on future climate forecasts (red = Seed Zone 1, yellow = Seed Zone 2, green = Seed Zone 3). The inset table clarifies the distance in kilometers between the present and predicted centroids for each seed zone.

We have ultimately determined the sample sizes for each seed zone to capture the majority of adaptive diversity existing in environments for the purpose of germplasm preservation (supplementary tables S8 and S9). To comprehensively capture the adaptive genomic diversity, our findings revealed that the greatest sampling demand was observed in SZ2, with 30.60 herbs required for 90% coverage of pan-adaptation loci, 58.70 herbs for 95%, and 280.80 herbs for 99%; similarly, for core adaptation loci, 16.10, 29.20, and 122.90 herbs were needed for 90%, 95%, and 99% coverage, respectively. Notably, despite the populations within SZ1 and SZ3 were identical, the datasets delineated distinct sampling requirements for each.

## Discussion

The goals of this study were to dissect the genomic basis underlying local adaptation within the *R. palmatum* complex, assess the genomic vulnerability of its populations in anticipated climatic conditions, define seed zones for effective germplasm preservation, and develop a quantitative framework to inform sampling strategies. To achieve these aims, we used SNPs arising from the whole-genome resequencing of 43 *R. palmatum* complex populations to identify signatures of environmental adaptation and to model the distribution of this adaptive variation across the landscape. Our comprehensive analyses revealed a polygenic basis for climate adaptation, which can be neatly summarized by the three distinct seed zones, from which we recommend collecting a relatively small number of individuals to effectively capture 95% of the wide adaptive diversity exhibited by our focal species. We emphasize the profound value of combining population genomic and landscape genomic approaches in predicting and monitoring species’ and populations’ responses to global climate change.

Characterizing GEAs remains a focal point of intense methodological and empirical advancement and there is a growing recognition of the relative contributions of climatic selection, geographic distance, and population structure to genetic differentiation within a given species (Savolainen, et al. 2013; Rellstab, et al. 2015; Lasky, et al. 2022). However, a significant challenge lies in the fact that these factors are often intertwined in natural landscapes (Wang, et al. 2013; Forester, et al. 2018). In our study focusing on the *R. palmatum* complex, we observed significant patterns of IBD and IBE based on both putatively neutral and adaptive SNPs. Furthermore, the results obtained through pRDA based on pan-adaptation and core adaptation loci revealed that a substantial portion of the explained variance in allele frequencies was shared among the neutral population structure, climate and geography. This likely reflects the post-glacial recolonization of our focal species, which created collinearity among its geographic distribution, genetic structure, and climatic gradients. This collinearity poses a challenge for genomic scans aiming to disentangle genetic variation arising from neutral and selective processes. Although it is common to conduct genome scans using proxies of neutral population structure to mitigate the risk of false positives (de Villemereuil, et al. 2014; Frichot, et al. 2015), such methods may obscure signals of climatic selection and hinder the identification of genes actually involved in climate adaptation (Anderson, et al. 2011; Savolainen, et al. 2013). In this study, we employed two GEA methods: LFMM, which treats population structure as a latent factor, and RDA without structure correction. We then utilized the pan-adaptation loci derived from the combined results of these two methods and the core adaptation loci identified through overlapping analysis to separately estimate genomic vulnerability, delineate seed zones, and evaluate germplasm conservation strategies. Although the dataset of core adaptation loci is thought to bolster confidence in frequency of true positives, it tends to skew detection towards regions undergoing pronounced selective sweeps (François, et al. 2016; Forester, et al. 2018). Conversely, the pan-adaptation loci, despite encompassing much more loci bearing weak selective signatures and those with small effects, may elevate the incidence of frequency of false positives. Nevertheless, a recent study has revealed that in scenarios where environmental gradients exhibit strong covariance with recolonization routes, we must acquiesce to a heightened rate of false positives if our aim is to pinpoint the genomic regions implicated in adapting to these gradients (Capblancq, et al. 2023).

Local adaptation typically involves a polygenic foundation governed by numerous genetic loci (Savolainen, et al. 2013; Barghi, et al. 2020). Under a polygenic adaptation model, natural selection operates to alter the allele frequencies of multiple causal loci (Barghi, et al. 2020; Fagny and Austerlitz 2021). Our findings suggest that a certain degree of multilocus selection—specifically, selection targeting numerous loci with small effects in response to environmental gradients—has influenced the adaptive capacity of populations across both western and eastern lineages. Within our focal species, adaptation manifests along two primary axes that distinguish populations in the western and eastern regions of its range. While the precise adaptive traits and their underlying genetic bases remain elusive in the *R. palmatum* complex, our discovery of climate-associated SNPs linked to a diverse array of genes and functional categories hints at the significant role of polygenic adaptation in the species’ evolutionary trajectory. Importantly, the putatively adaptive SNPs identified in this study await validation with phenotypic data, and their molecular functions remain unknown. Furthermore, considering that the genomic diversity of our target species is significantly lower compared to other perennial herbs such as alfalfa (Zhang, et al. 2024) and lettuce (Wei, et al. 2021), epigenetic variations, including DNA methylation, can emerge rapidly in species with limited genetic diversity as a response to environmental stimuli and facilitate swift adaptation to novel environments (Huang, et al. 2017; Aagaard, et al. 2022). Future research endeavors, harnessing advanced analytical techniques from functional genomics and incorporating epigenomic variations into genomic offset estimations, hold the potential to enhance prediction accuracy. Such research can offer a more mechanistic insight into climate (mal)adaptation and provide direct evidence elucidating the functional roles of these outlier loci.

Exploring and understanding the evolutionary forces and genomic architecture that underlie contemporary climate adaptation offer a pivotal foundation for predicting species’ responses to ongoing climatic shifts (Aguirre-Liguori, et al. 2021; Bernatchez, et al. 2024). In light of the swift pace of climate change, adaptation necessarily hinges on preexisting genetic variation, such as standing genetic variation, both within and across populations (Aitken, et al. 2008; Barrett and Schluter 2008; Bomblies and Peichel 2022). Our species-level estimations of genomic offsets, derived from pan-adaptation and core adaptation loci under various climatic scenarios, generally revealed that populations belonging to the eastern lineage were likely to encounter more pronounced maladaptation. Conversely, populations occupying the western portion of the range were projected to experience a more tempered maladaptation. Populations devoid of preexisting adaptive variability may remain unable to acclimatize in the forthcoming decades without the influx of adaptive variation from external populations. To alleviate local maladaptation, assisted gene flow from populations exhibiting lower genomic offsets within the same lineage emerges as a potential strategy. This is particularly pertinent given the limited gene flow between western and eastern lineages and the potential risk of outbreeding depression resulting from the admixture of individuals from distinct evolutionary lineages (Hufford and Mazer 2003; Mijangos, et al. 2015; Grummer, et al. 2022). The feasibility of this resilience strategy is further supported by the inconsistent genomic offsets observed in species with high genetic structure, as evident in both our study and other recent investigations (Lind, et al. 2024; Lind and Lotterhos 2024). Therefore, future assessments of genomic offset should be conducted at the lineage level for species with high genetic structure, if the populations and individuals per population across different lineages meet the criteria of sampling density (Aguirre-Liguori, et al. 2023). Although certain studies have emphasized the need for caution when utilizing genomic offset estimates to inform management decisions, due to inherent concerns such as the presence of local adaptation (Lind and Lotterhos 2024), fixed GEAs across varying climates (Capblancq, et al. 2020; Rellstab 2021), disregard for phenotypic plasticity (Foden, et al. 2019) and genetic load (Brady, et al. 2019), which can collectively affect the reliability of genomic offset; nevertheless, for immediate assisted gene flow efforts, offset models may offer the highest degree of precision and relevance.

An effective delineation of seed zones must accurately reflect patterns of local adaptation among populations of the targeted species, ensuring that germplasms sourced from each zone can be strategically deployed within environmentally similar ranges to optimize adaptation and productivity, while maintaining acceptable risk levels (Li, et al. 2017). Historically, characterizing the nature and extent of local adaptation, as well as the subsequent delineation of seed zones, has typically relied on reciprocal common garden experiments (Bower, et al. 2014; Bucharova, et al. 2017). However, given the impossibility of acquiring wild seeds representative of all, or even most, historically occupied climates by the species, we have sought to utilize correlations between multivariate genotypes and their respective climate spaces as a proxy for adaptation. In this study, by leveraging adaptive genomic and climate data specific to our focal species, we elucidated that the range of the *R. palmatum* complex could be most parsimoniously partitioned into three distinct seed zones, which best represent the species’ adaptive boundaries. Notably, the longitudinal boundaries of these three seed zones roughly correspond to the phylogeographical divide between western and eastern lineages (Wang, et al. 2018; Feng, et al. 2020), which have traditionally relied on genetic variation to delineate management units (Funk, et al. 2012; García-Dorado and Caballero 2021). This concordance suggests a parallelism between population structure and adaptation, particularly when axes of recolonization align with those of climate differentiation (Sandercock, et al. 2024).

We also estimated the sampled individuals for each seed zone based on the adaptive genomic diversity derived from pan-adaptation and core adaptation loci, respectively. The differing estimations between these two datasets highlight the need for differentiated sampling strategies. The divergent sampling strategies imply that, when formulating germplasm preservation strategies, relying on sampling recommendations based on pan-adaptation loci is preferable, as it enables the preservation of maximal adaptive genetic diversity, thereby enhancing future population resilience, albeit with higher false-positive rates compared to core adaptation loci. However, given the possibility that some adaptive genomic diversities may remain undetected even with the utilization of pan-adaptation loci in our investigation, it would be wise to adopt a cautious approach and conduct extensive sampling within the three seed zones when formulating a restoration plan for our focal species.

## Methods

### Sampling, sequencing and SNP calling

In line with the previous recommendations for landscape genomic studies (Bragg, et al. 2015; Aguirre-Liguori, et al. 2023), we collected 213 accessions of the *R. palmatum* complex from 43 natural populations throughout its distribution range for whole-genome resequencing. Genomic DNA was extracted from fresh leaves or roots and subsequently sequenced on the DNBSEQ-T7 platform, targeting a minimum coverage depth of ≥ 15× per individual. Following the manufacturer’s protocols, DNA libraries were crafted and subjected to 150 base pair (bp) paired-end sequencing.

Raw sequence data underwent rigorous quality control using Trimmomatic v0.39 (Bolger, et al. 2014), eliminating adapter sequences and low-quality bases (Phred score < 30). The high-quality reads were then mapped to the *R. palmatum* genome (Zhang, et al. 2024) using BWA v0.7.17 (Li and Durbin 2009) with default parameters. The resulting SAM files were transformed into sorted BAM files and indexed utilizing SAMtools v0.1.19 (Li, et al. 2009). PCR duplicates were marked with Picard (https://github.com/broadinstitute/picard), and candidate variants were identified on a per-chromosome basis employing the HaplotypeCaller from GATK v4.2.6.1 (McKenna, et al. 2010). These individual GVCF files were amalgamated into a single GVCF per sample using GatherVcfs, which were further processed by GenotypeGVCFs to yield a raw variant call file (VCF). Quality filtration of this VCF was performed with the VariantFiltration module within GATK, applying stringent criteria (QD < 2.0 || FS > 60.0 || MQRankSum < −12.5 || ReadPosRankSum < −8.0 || SOR > 3.0 || MQ < 40.0). Lastly, BCFtools (Danecek, et al. 2011) was employed to remove INDELs and multiallelic SNPs under default settings, leading to a refined dataset encompassing 14,299,652 SNPs across the 213 accessions.

Next, strict criteria for soft filtration (Marees, et al. 2018) were implemented to further curtail the quantity of low-quality genotype calls via PLINK v1.9 (Purcell, et al. 2007). The parameters employed for SNP filtering in PLINK encompassed a minor allele frequency threshold of 0.05 (--maf 0.05), a maximum permissible per-SNP missingness of 2% (--geno 0.02), and a maximum allowed per-sample missingness of 2% (--mind 0.02). This filtering process resulted in the identification of 1,432,892 high-quality SNPs from 201 samples. Additionally, SNPs exhibiting high correlation were eliminated through LD-based SNP pruning in PLINK, utilizing the parameter “--indep-pairwise 50 10 0.2” (this removed any SNP having a correlation coefficient (*r*^2^) exceeding 0.2 with another SNP within sliding windows containing 50 SNPs, with a step advancement of 10 SNPs across the genome). The resultant SNP dataset, comprising 378,620 SNPs from 201 accessions, was subsequently utilized for clustering analyses.

### Genetic diversity, linkage disequilibrium and population structure analyses

We employed the R package *hierfstat* (Goudet 2005) to ascertain population differentiation (*F*_ST_) (Weir and Cockerham 1984) amongst various populations, and further assessed diversity indices, specifically observed heterozygosity (*H*_O_) and genetic diversity (*H*_S_) within these populations. Additionally, we utilized PLINK to determine nucleotide diversity (π) within our focal species’ populations. Tajima’s *D* statistics were also calculated for the all populations of our focal species utilizing VCFtools v0.1.15 (Danecek, et al. 2011) in 100-kbp non-overlapping windows. To further evaluate and compare the patterns of linkage disequilibrium among distinct population groups, PopLDdecay v.3.40 (Zhang, et al. 2018) was employed to calculate the squared correlation coefficient (*r*^2^) between SNP pairs with a MAF > 0.05 within a 100-kbp window, subsequently averaging these values across the entire genome.

In our study, three methodologies were utilized to ascertain the genetic structure of the focal species. Initially, a PCA was executed in R v4.2.1 (Team 2013) to assess the genetic structure, preceded by the acquisition of eigenvalues from 378,620 SNPs via PLINK. Secondly, a DAPC was conducted to determine the number of discrete clusters, facilitated by the R package *adegenet* (Jombart and Ahmed 2011). Lastly, STRUCTURE v2.3.4 (Pritchard, et al. 2000) was employed to investigate the population structure across all individuals, wherein the predefined number of genetic clusters *K*, representing population ancestries, ranged from 1 to 10 for both the entire population and those belonging to western lineages (see results). The optimal *K* was deduced through likelihood plots (Pritchard, et al. 2000) and the Δ*K* method (Evanno, et al. 2005), implemented in the web-based software, STRUCTURE HARVESTER (Earl and vonHoldt 2012). The programs CLUMPP v1.1.2 (Jakobsson and Rosenberg 2007) and DISTRUCT v1.1 (Rosenberg 2004) were subsequently utilized to consolidate the individual STRUCTURE outcomes.

### Estimation of demographic history

Given the recent divergence of western and eastern lineage populations of our focal species (Wang, et al. 2018; Feng, et al. 2020), we estimated the demographic histories of the three genetic clusters separately utilizing the stairway plot v2 (Liu and Fu 2015) and SMC++ v1.15.4 (Terhorst, et al. 2017). Assuming a mutation rate of 7.0×10^−9^ (per site per generation) and adhering to a generation time of 5 years (Feng, et al. 2020), the scaled outcomes were subsequently transformed into years and effective population size (*Ne*). Median estimates and confidence intervals for the *Ne* in the stairway plot were calculated using the built-in bootstrap function with 200 subsets of the input data. To infer the demographic history with SMC++, we converted a VCF file containing high-quality SNPs to the SMC format using the vcf2smc command. Subsequently, the estimate command was executed for each group using default parameters.

### Determining climate-associated SNPs and their impact on genetic variation

In our study, we employed two distinct methods to pinpoint SNPs that exhibit strong associations with BIOCLIM variables across the genomic regions of our focal species. To detect GEAs using the LFMM model implemented within the R package *LEA* (Frichot and Francois 2015), we executed five independent Markov chain Monte Carlo runs, incorporating a latent factor of K[=[3 to account for population structure. The *P*-values obtained from all five runs were subsequently aggregated and adjusted for multiple testing using a false discovery rate correction, with a threshold of *P*[<[0.01. Due to the fact that LFMM fails to account for the correlation structure among environmental variables, we adhered to the developers’ guidelines and condensed the six climate variables (see below) into synthetic variables based on principal component axes. We contend that this decision is justified, given that climate variables, whether utilized in the analysis or excluded, exhibit correlations with one another within each specific context, thereby complicating our endeavor to causally ascribe associations to any particular variable.

RDA has emerged as one of the most effective multivariate GEA approaches, demonstrating low false-positive rates across various demographic scenarios (Forester, et al. 2016; Forester, et al. 2018). After considering the ranked importance of the 19 BIOCLIM variables, which were assessed using GF analyses in the R package *gradientForest* (Ellis, et al. 2012), along with the correlations among these variables, we selected six BIOCLIM variables (specifically, BIO2, BIO3, BIO4, BIO10, BIO12, and BIO15) with pairwise correlation coefficients |*r*| < 0.8 for the RDA, implemented in the R package *vegan* (Oksanen, et al. 2019). SNPs were regarded as putatively adaptive loci if their loading values exceeded 2.7 standard deviations from the mean loading values. We designated the outlier loci that were identified by both approaches as core adaptation loci, whereas those detected by either method were classified as pan-adaptation loci. Following this classification, we employed BEDTools v2.31.0 (Quinlan and Hall 2010) to annotate both core adaptation and pan-adaptation loci onto their corresponding genes. Subsequently, we performed a gene ontology (GO) enrichment analysis on the annotated genes using the R package *clusterProfile* (Wu, et al. 2021), which enables identifying the significantly enriched biological pathways associated with these genes.

In addition, we performed Mantel and partial Mantel tests to ascertain the influence of IBD and IBE on shaping the spatial genetic variation of both adaptive and neutral loci. The significance of associations between *F*_ST_ (*F*_ST_/1 − *F*_ST_) and geographic as well as environmental distances (after accounting for the effect of geographic distance) was determined through 999 permutations using the R package *vegan*. Furthermore, we conducted pRDA to estimate the relative contributions of geography, neutral population structure, and environment in driving genetic variation across the *R. palmatum* complex populations. To construct one full model and three partial models, we utilized three datasets: (i) six selected BIOCLIM variables (‘clim’); (ii) loadings from the first three principal components of a genetic PCA based on neutral loci (‘struct’); and (iii) population coordinates, including latitude and longitude (’geog’). These models were built using the R package *vegan*. Population allele frequencies served as the response variable in all four models, and the significance of the explanatory variables was evaluated using 999 permutations with the ‘anova.cca’ function in the *vegan* package.

### Genomic offset estimation

For each sampling location, we obtained the 19 BIOCLIM variables corresponding to the current (1970-2000) and future (2050 and 2090) timeframes at a 2.5-min resolution from WorldClim v2.1 (Fick and Hijmans 2017). Both future datasets encompass four distinct climate models: BCC-CSM2-MR, ACCESS-CM2, MIROC6, and MPI-ESM1-2-HR, along with two common socioeconomic pathways (SSPs), namely SSP126 and SSP585. we integrated the four future climate models across the same time scales and SSPs via the R package *raster* (Hijmans, et al. 2015). To assess the genomic offset of our focal species in response to the anticipated climate changes, we employed two distinct methodologies.

Firstly, we utilized a nonparametric, machine-learning GF model to evaluate the genomic offset across the species range of *R. palmatum* complex. This was achieved through the utilization of the R package *gradientForest*, following the methodologies outlined by Fitzpatrick and Keller (2015). For each climate model considered, we constructed a GF model to estimate the genomic offset under varying climatic conditions, incorporating six unrelated environmental variables. The GF model predicts genomic offset as the divergence in climatically adaptive SNPs between current and future climates within the same location, and elevated values of genomic offset serve as indicators of heightened risk of maladaptation under forecasted climatic conditions (Fitzpatrick and Keller 2015). When constructing GF model, we utilized 500 regression trees per SNP, set a variable correlation threshold of 0.5, and maintained the default settings for all other parameters. In addition, after considering recent studies that focus on evaluating the efficiencies of genomic offset in predicting maladaptation to future climates (Aguirre-Liguori, et al. 2023; Lind, et al. 2024), we calculated and contrasted these offsets between two distinct types of datasets (i.e., the datasets of core adaptation loci and pan-adaptation loci) at both the species and lineage levels.

In addition, as a complementary approach to GF, we investigated the influence of adaptive variation across diverse landscapes adopting the methodology of Capblancq et al (2020). Initially, we conducted a secondary RDA utilizing outliers previously identified through LFMM and RDA methods. These outliers constitute an adaptively enriched genetic space (Steane, et al. 2014), and performing a secondary RDA on these specific loci enables the identification of environmental variables most strongly correlated with putatively adaptive variation. Subsequently, we constructed composite indices using scores from the six uncorrelated environmental variables along the first two RDA axes, aiming to predict the adaptive score of individuals within their environment. The GEAs employed here can be extrapolated to future climates, allowing for predictions of potential shifts in adaptive optima triggered by climatic changes. We replicated the procedure used for current climatic conditions to forecast the optimal adaptive index based on future climate scenarios (specifically, 2050 and 2090) at two distinct SSPs. The Euclidean distance between the adaptive indices under present and future climates for each pixel offers an estimation of the genomic offset (Capblancq and Forester 2021). We aggregated the genomic offsets across both RDA axes and visualized the results with the aid of R packages *raster* and *ggplot2* (Wickham 2011). Akin to the genomic offset evaluated using the GF model, the genetic offset was computed among datasets containing two types of outliers, both at the species and lineage levels.

Finally, in order to evaluate the genetic load within the *R. palmatum* complex population, genetic variants were annotated as either synonymous or non-synonymous. Understanding genetic load is important to assess its impact on the health and viability of endangered populations, because it affects the extinction risk and recovery potential of populations (Bertorelle, et al. 2022; Dussex, et al. 2023). For the non-synonymous variants, SIFT scores were computed utilizing the SIFT 4G software (Vaser, et al. 2016), with UniRef100 serving as the protein database. Subsequently, these variants were categorized into four distinct groups: loss of function (LOF), deleterious (with a SIFT score < 0.05), tolerated (with a SIFT score ≥ 0.05), or synonymous. Furthermore, the determination of the derived versus ancestral allelic state at each SNP position was carried out by employing est-sfs (Keightley and Jackson 2018), with *Rheum nobile* (Feng, et al. 2023) serving as the outgroup. The genetic load was approximated by calculating the ratio of the number of derived mutations (comprising LOF and deleterious mutations) to the number of synonymous variants. In addition, we employed the ‘stat_cor’ function in the *ggpubr* package (Kassambara 2018) to assess the correlation between the genomic offsets under two future scenarios (SSP126 and SSP585) across two distinct timescales (2050 and 2090) and genetic diversity across all populations, as well as the association between genomic offset and proxies of genetic load across all populations.

### Delineating seed zones and determining their changes under future climates

In the context of multivariate adaptive genomic variation, seed zones represent areas of relative homogeneity, serving as a basis for the *ex situ* germplasm conservation and guiding the breeding of climate-matched restoration populations (Sandercock, et al. 2024). Following the method of Yu *et al*. (Yu, et al. 2022), we employed the partitioning around medoids (k-medoids) algorithm^111^ along with the R package *factoextra* (Kassambara and Mundt 2017) to assess the variation within clusters for various cluster numbers. This assessment was based on GF-predicted continuous genomic variation utilizing two distinct outlier datasets. We investigated cluster numbers ranging from two to ten, determining the optimal count by identifying the elbow point–the juncture where adding an additional cluster results in significantly less reduction of within-cluster variation compared to previous additions. The chosen number of clusters was then utilized to categorize the spatial distribution of GF-predicted continuous genomic variation into GF-based seed zones across the ranges of our focal species.

To anticipate the shifts in defined seed zones from the present to the future climates, we followed the approach of Sandercock *et al*. (2024). Utilizing the ‘knn.dist’ function from the R package *FNN* (Beygelzimer, et al. 2013) with k = 1 (representing the number of nearest neighbors to search), we matched each present-day location (pixel) with its closest future analog based on the minimal genomic offset. This approach facilitates effective prediction of the minimal migration path required to mitigate the climate maladaptation. For each seed zone, we calculated the centroids separately for pixels representing the current climate conditions and those uniquely aligned with future scenarios. Subsequently, we quantified the migratory distances between these present and future centroids, thereby deriving an estimation of the average minimum migration distance required for each seed zone.

### Evaluating the sampling numbers for *ex situ* germplasm conservation

To assess the requisite sampling sizes for each seed zone, aimed at recapitulating genomic variation at adaptive loci and guiding germplasm conservation strategies, we employed linear regression analysis to compare allele frequencies of putatively adaptive loci in the entire population with those from bootstrap samples of varying sizes (Sandercock, et al. 2024). The Python script facilitating this process is accessible at: https://github.com/alex-sandercock/Capturing_genomic_diversity. Employing the R² values derived from these regressions as a proxy for the percentage of diversity captured in each iteration, we iteratively augmented the sample size until the R² achieved or surpassed a predefined threshold (e.g., 90%). This procedure was replicated 100 times, and the recommended sampling numbers for each designated seed zone were averaged across all iterations. The outcome represents an estimation of the mean sampling number required for each seed zone to achieve the desired coefficient of determination between the sample’s adaptive genomic composition and that of the full dataset.

## Acknowledgements

We extend our heartfelt gratitude to Prof. Fang Du from Beijing Forestry University, Prof. Nian Wang from Shandong Agricultural University, and Dr. Shan-Shan Zhu from Ningbo University for their invaluable suggestions and comments, which have significantly enhanced the initial manuscript. This research was funded by the National Natural Science Foundation of China (32271550 to L.F., 32371911 to Y.W., and 32370408 to X.W.) and the Shaanxi Biochemical Basic Science Research Project (22JHZ005 to X.W.).

## Author contributions

L.F. conceived and supervised the study. T.Z., Z.W., and X.W. performed the sample collections. L.F., C.W., L.Z., and Y.W. conducted all bioinformatics analyses. L.F. and C.W. wrote the manuscript, with contributions from J.W. and X.W. All authors approved the final manuscript.

## Data availability

The whole-genome resequencing data for 213 individuals belonging to the *Rheum palmatum* complex have been deposited in the National Genomics Data Center (https://ngdc.cncb.ac.cn) under the accession number PRJCA033457. The VCF dataset, alongside the location records of 43 *R. palmatum* complex populations and the climatic data utilized for the identification of putatively adaptive loci and the estimation of genomic offset, are accessible at https://doi.org/10.6084/m9.figshare.28060457.v1. All scripts used in this study are available at https://github.com/Feng-Li-hub/Landscape-genomes-of-Rheum-palmatum-complex.

## Supplementary information

### Supplementary figures

**Supplementary fig. S1** Population assignment outcomes for 201 individuals within the *Rheum palmatum* complex. Panels (a) and (c) depict the mean posterior probability of the dataset across *K* = 1-10 (five replicates) for the entire set of 43 populations and the western lineage populations, respectively, with the standard deviation of each mean Ln(*K*) value provided. Panels (b) and (d) present the Δ*K* values associated with varying *K* values for the total 43 populations and the western lineage populations, respectively. Panels (e) and (f) illustrate the population assignment results for the 201 *Rheum palmatum* complex individuals based on principal component analysis (PCA) and discriminant analysis of principal components (DAPC), respectively.

**Supplementary fig. S2** Population assignment outcomes for 201 individuals of the *Rheum palmatum* complex based on pan-adaptation loci (a, b) and core adaptation loci (c, d). Panels (a) and (c) depict clustering results obtained via principal component analysis (PCA), whereas panels (b) and (d) show clustering results from discriminant analysis of principal components (DAPC).

**Supplementary fig. S3** Demographic histories of the three lineages within the *Rheum palmatum* complex. Panels (a) and (b) depict the inferred changes in population effective size for the Eastern, WE1, and WE2 lineages using SMC++ and stairway plot methods, respectively. The dashed colored lines in (b) represent the 95% confidence intervals for the changes in population effective size. (c) Linkage disequilibrium (LD) decay estimated by PopLDdecay for the three groups.

**Supplementary fig. S4** Spearman’s correlation coefficient (two-sided test) among 19 environmental variables (positioned above the diagonal), alongside the ranked importance of these variables determined through gradient forest analysis (displayed below the diagonal). Variables marked with an asterisk (*) have been chosen for further genotype-environment association analysis.

**Supplementary fig. S5** (a) Venn diagram illustrates putatively adaptive SNPs identified within the *Rheum palmatum* complex through latent factor mixed model (LFMM) and redundancy analysis (RDA). (b) Distribution of annotated genes across the whole-genome of *Rheum palmatum*, with adaptive SNP annotation genes corresponding to pan-adaptation loci (purple circles) and core adaptation loci (orange triangles).

**Supplementary fig. S6** Patterns of genetic differentiation within the *Rheum palmatum* complex. Panels (a) and (c) represent the isolation-by-distance analysis (two-sided Mantel test) for (a) pan adaptation loci and (c) core adaptation. Shadows represent the 95% confidence interval of linear regression. Panels (b) and (d) represent the isolation-by-environment analysis (two-sided partial Mantel test) for (b) pan-adaptation loci and (d) core adaptation loci, with shadows indicating the 95% confidence interval.

**Supplementary fig. S7** The adaptive landscape of the *Rheum palmatum* complex. (a) The adaptively enriched genetic space showing association between pan-adaptation loci and the six key climatic drivers of adaptation. (b) Spatial projection of adaptive genetic turnover across the potential range of *Rheum palmatum* complex.

**Supplementary fig. S8** The adaptive landscape of the *Rheum palmatum* complex. (a) The adaptively enriched genetic space showing association between core adaptation loci and the six key climatic drivers of adaptation. (b) Spatial projection of adaptive genetic turnover across the potential range of *Rheum palmatum* complex.

**Supplementary fig. S9** Predicted genomic offsets across the range of *Rheum palmatum* complex in response to future climates, derived from core adaptation loci using gradient forest (panels a-d) and redundancy analysis (panels e-h). Panels (a) and (e) represent the scenarios corresponding to the shared socioeconomic pathways SSP126 in 2050, while (b) and (f) depict SSP126 in 2090. Similarly, (c) and (g) illustrate SSP 585 in 2050, and (d) and (h) show SSP585 in 2090. The size of the blue circles in each panel corresponds to the magnitude of the genomic offset under the respective climate scenarios.

**Supplementary fig. S10** Predicted genomic offsets of the western and eastern lineage populations derived from species-level estimations, utilizing both pan-adaption loci (panels a and b) and core adaptation loci (panels c and d) via redundancy analysis. On the x-axis, the numerals 5026 and 9026 denote the scenarios corresponding to the shared socioeconomic pathways SSP126 in 2050 and 2090, respectively, while 5085 and 9085 represent the SSP585 scenarios in 2050 and 2090, respectively.

**Supplementary fig. S11** Consistency in environmental accuracy (depicted by light purple circles) and weighted importance rankings (represented by light green rectangles) as outputted by trained gradient forest models utilizing pan-adaptation loci and core adaptation loci at species and lineage levels.

**Supplementary fig. S12** Predicted genomic offsets at the lineage level based on pan-adaptation loci (panels a-h) and core adaptation loci (panels i-p) using redundancy analysis. Panels (a) and (i) depict the offsets for western lineage populations under the scenarios corresponding to the shared socioeconomic pathways SSP126 in 2050, while (b) and (j) represent the same for 2090. Panels (c) and (k) exhibit offsets for SSP585 in 2050, and (d) and (l) for 2090. For eastern lineage populations, panels (e) and (m) display offsets under SSP126 in 2050, (f) and (n) for 2090, (g) and (o) for SSP585 in 2050, and finally, (h) and (p) for 2090.

**Supplementary fig. S13** Relationships between various proxies of genetic load (y-axis) and genomic offset (x-axis) via gradient forest based on pan-adaptation loci under SSP126 socioeconomic pathway scenarios for 2050 (a) and 2090 (b), and SSP585 scenarios for 2050 (c) and 2090 (d). (a) Relationship between genomic offset and the ratio of derived deleterious variants to derived synonymous variants. (b) Relationship between genomic offset and the ratio of derived tolerant variants to derived synonymous variants. (c) Relationship between genomic offset and the ratio of loss-of-function variants to derived synonymous variants. (d) Relationship between genomic offset and nucleotide diversity.

**Supplementary fig. S14** Relationships between various proxies of genetic load (y-axis) and genomic offset (x-axis) via redundancy analysis based on pan-adaptation loci under SSP126 socioeconomic pathway scenarios for 2050 (a) and 2090 (b), and SSP585 scenarios for 2050 (c) and 2090 (d). (a) Relationship between genomic offset and the ratio of derived deleterious variants to derived synonymous variants. (b) Relationship between genomic offset and the ratio of derived tolerant variants to derived synonymous variants. (c) Relationship between genomic offset and the ratio of loss-of-function variants to derived synonymous variants. (d) Relationship between genomic offset and nucleotide diversity.

**Supplementary fig. S15** Relationships between various proxies of genetic load (y-axis) and genomic offset (x-axis) via gradient forest based on core adaptation loci under SSP126 socioeconomic pathway scenarios for 2050 (a) and 2090 (b), and SSP585 scenarios for 2050 (c) and 2090 (d). (a) Relationship between genomic offset and the ratio of derived deleterious variants to derived synonymous variants. (b) Relationship between genomic offset and the ratio of derived tolerant variants to derived synonymous variants. (c) Relationship between genomic offset and the ratio of loss-of-function variants to derived synonymous variants. (d) Relationship between genomic offset and nucleotide diversity.

**Supplementary fig. S16** Relationships between various proxies of genetic load (y-axis) and genomic offset (x-axis) via redundancy analysis based on core adaptation loci under SSP126 socioeconomic pathway scenarios for 2050 (a) and 2090 (b), and SSP585 scenarios for 2050 (c) and 2090 (d). (a) Relationship between genomic offset and the ratio of derived deleterious variants to derived synonymous variants. (b) Relationship between genomic offset and the ratio of derived tolerant variants to derived synonymous variants. (c) Relationship between genomic offset and the ratio of loss-of-function variants to derived synonymous variants. (d) Relationship between genomic offset and nucleotide diversity.

**Supplementary fig. S17** Intra-cluster variation (black solid line) and its reduction (blue bars) with increasing number of clusters, as estimated from continuous genomic variation predicted by gradient forest based on pan-adaptation loci (a) and core adaptation loci (b) across the range of *Rheum palmatum* complex.

**Supplementary fig. S18** Predicted seed zones based on core adaptation loci and their range shifts to minimize genomic offset under climate change. Panel (a) represents predicted three seed zones for the *Rheum palmatum* complex using gradient forest (GF). The biplot in subfigure illustrates the GF-predicted genomic variation used for seed zone delineation. Panels (b-e) represent the predicted range adjustments for each seed zone to minimize genomic disparities derived from pan-adaptation loci under various climate scenarios. Solid-colored numerals denote the current centroids of the seed zones, whereas numerals on a white background indicate the anticipated centroid shifts in alignment with future climate projections. Panels (b) and (c) correspond to the SSP126 socioeconomic pathway scenarios in 2050 and 2090, respectively. Panels (d) and (e) reflect the SSP585 scenarios in 2050 and 2090, respectively. The colored pixels represent the predicted relocation points for each seed zone, where genomic disparities are minimized based on future climate forecasts (red = Seed Zone 1, yellow = Seed Zone 2, green = Seed Zone 3). The inset table clarifies the distance in kilometers between the present and predicted centroids for each seed zone.

### Supplementary tables

**Supplementary table S1**. Geographical sampling information and summary statistics of genetic diversity and Tajima’s *D* values across populations in this study.

**Supplementary table S2**. Summary statistics of whole genome resequencing data for samples used in this study.

**Supplementary table S3**. Population structure (pairwise *F*_ST_) among the 43 populations of the *Rheum palmatum* complex.

**Supplementary table S4**. Environmental variables used in this study derived from WorldClim.

**Supplementary table S5**. List of the genes associated with the putatively adaptive loci identified by genomic scans, including gene names, a description of the associated protein functions.

**Supplementary table S6**. The influence of climate, geography and neutral genetic (excluding the pan-adaptation loci) structure on genetic variation of the *Rheum palmatum* complex decomposed with partial redundancy analysis (pRDA) using pan-adaptation loci.

**Supplementary table S7**. The influence of climate, geography and neutral genetic (excluding the pan-adaptation loci) structure on genetic variation of the *Rheum palmatum* complex decomposed with partial redundancy analysis (pRDA) using core adaptation loci.

**Supplementary table S8**. Sampling estimates for capturing adaptive diversity across wild seed zones of the *Rheum palmatum* complex based on pan-adaptation loci.

**Supplementary table S9**. Sampling estimates for capturing adaptive diversity across wild seed zones of the *Rheum palmatum* complex based on core adaptation loci.

