## Supporting figure S1-S18 for "Harnessing landscape genomics to evaluate genomic vulnerability and future climate resilience in an East Asia perennial"

**Supplementary figures**

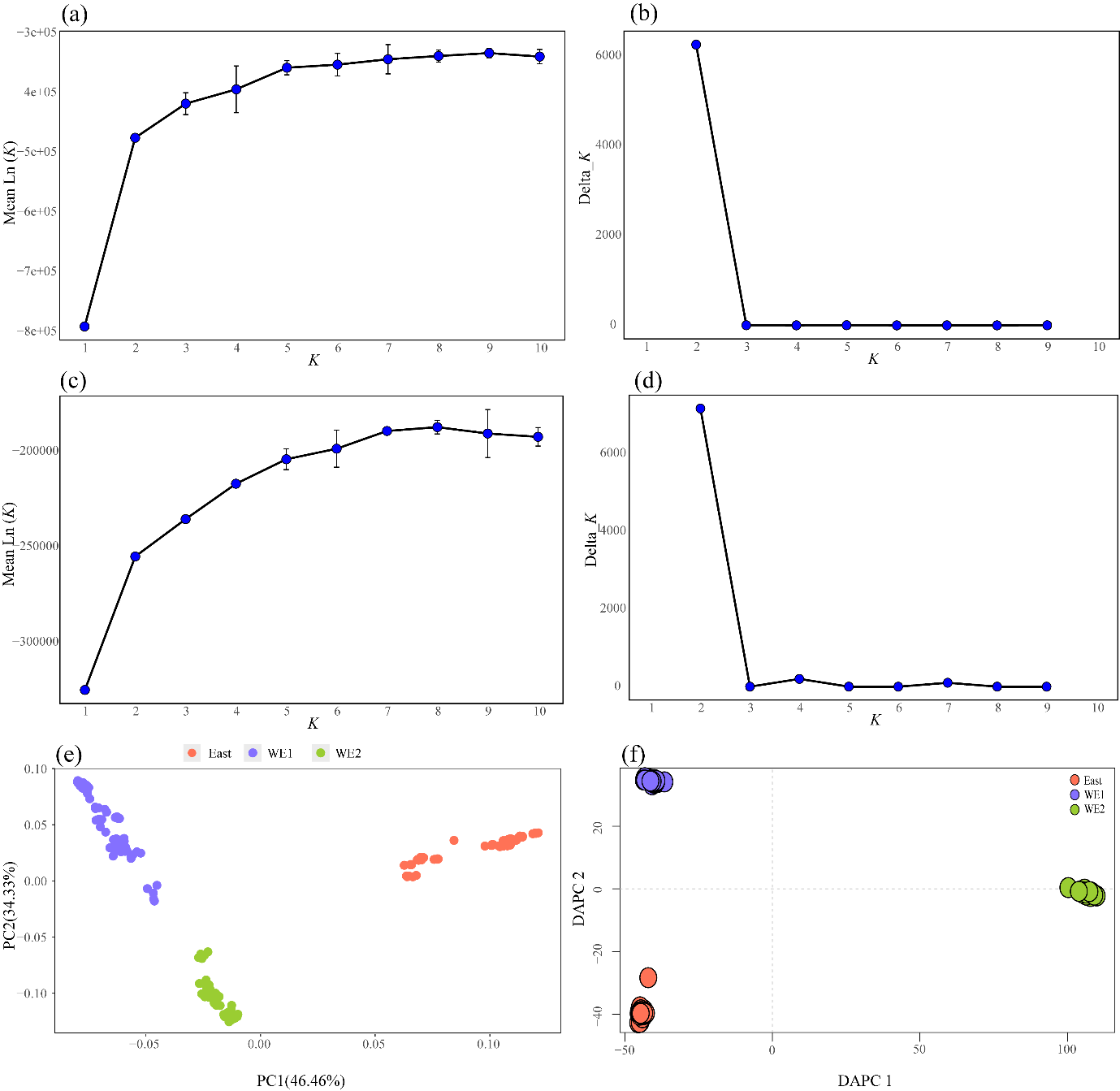

**Supplementary fig. S1** Population assignment outcomes for 201 individuals within the *Rheum palmatum* complex. Panels (a) and (c) depict the mean posterior probability of the dataset across *K* = 1-10 (five replicates) for the entire set of 43 populations and the western lineage populations, respectively, with the standard deviation of each mean Ln(*K*) value provided. Panels (b) and (d) present the *∆*K values associated with varying *K* values for the total 43 populations and the western lineage populations, respectively. Panels (e) and (f) illustrate the population assignment results for the 201 *Rheum palmatum* complex individuals based on principal component analysis (PCA) and discriminant analysis of principal components (DAPC), respectively.

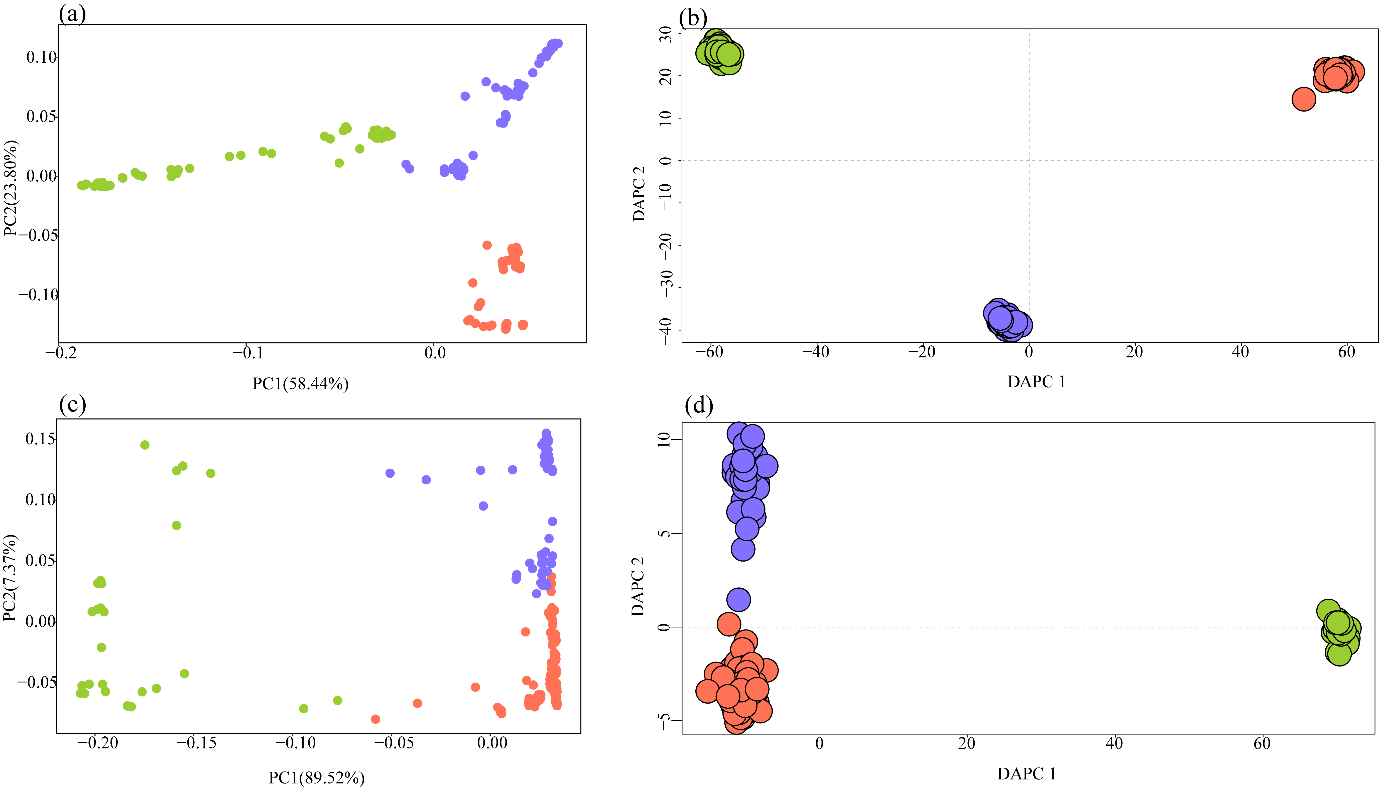

**Supplementary fig. S2** Population assignment outcomes for 201 individuals of the *Rheum palmatum* complex based on pan-adaptation loci (a, b) and core adaptation loci (c, d). Panels (a) and (c) depict clustering results obtained via principal component analysis (PCA), whereas panels (b) and (d) show clustering results from discriminant analysis of principal components (DAPC).

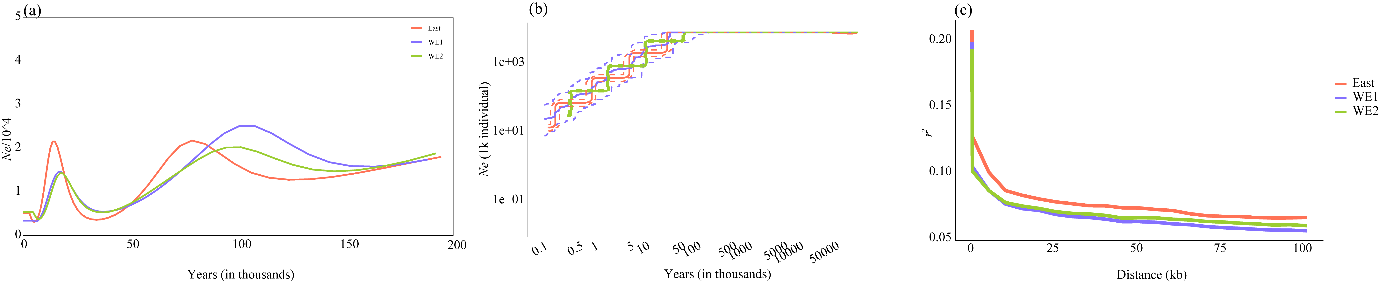

**Supplementary fig. S3** Demographic histories of the three lineages within the *Rheum palmatum* complex. Panels (a) and (b) depict the inferred changes in population effective size for the Eastern, WE1, and WE2 lineages using SMC++ and stairway plot methods, respectively. The dashed colored lines in (b) represent the 95% confidence intervals for the changes in population effective size. (c) Linkage disequilibrium (LD) decay estimated by PopLDdecay for the three groups.

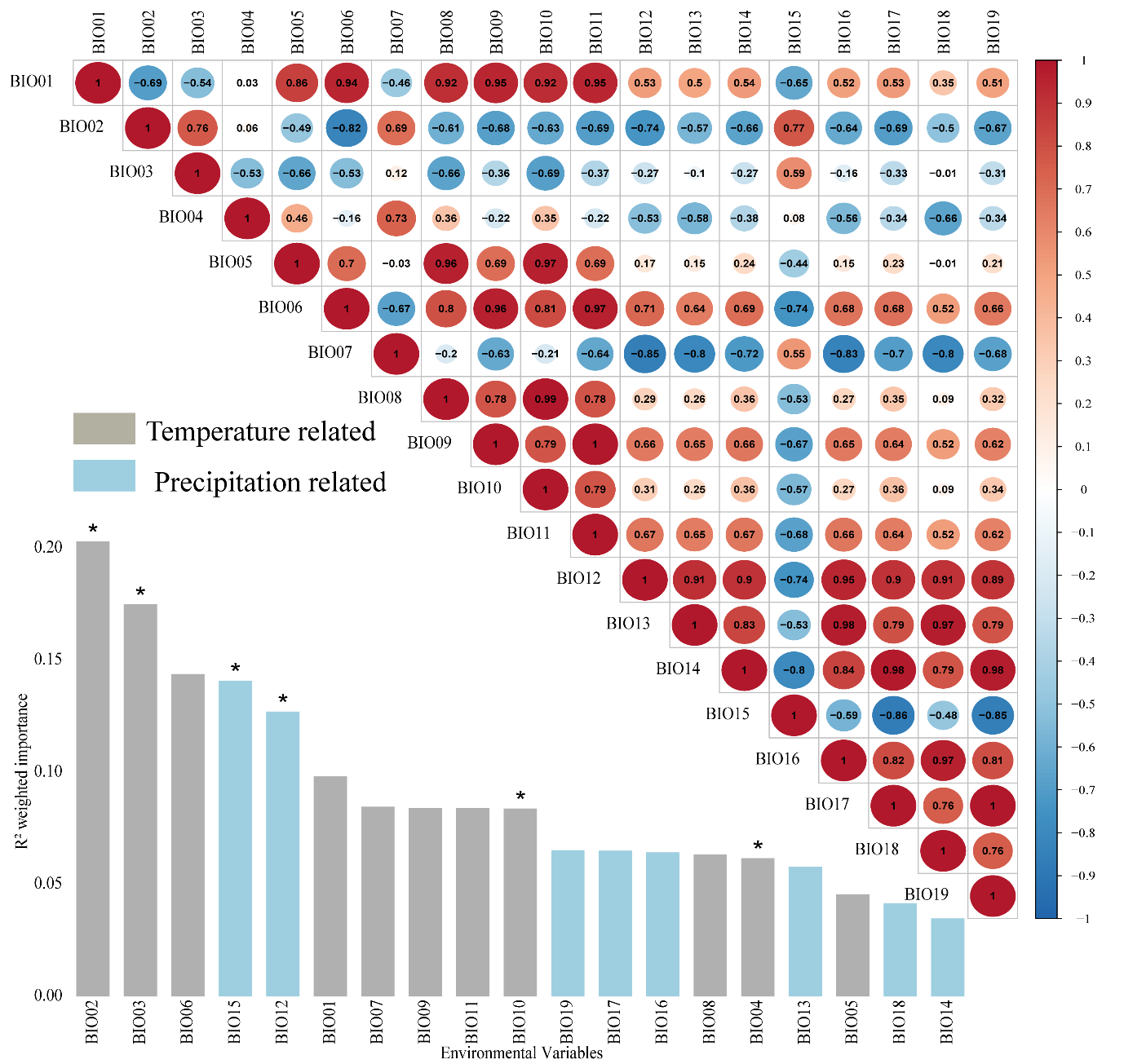

**Supplementary fig. S4** Spearman's correlation coefficient (two-sided test) among 19 environmental variables (positioned above the diagonal), alongside the ranked importance of these variables determined through gradient forest analysis (displayed below the diagonal). Variables marked with an asterisk (*) have been chosen for further genotype-environment association analysis.

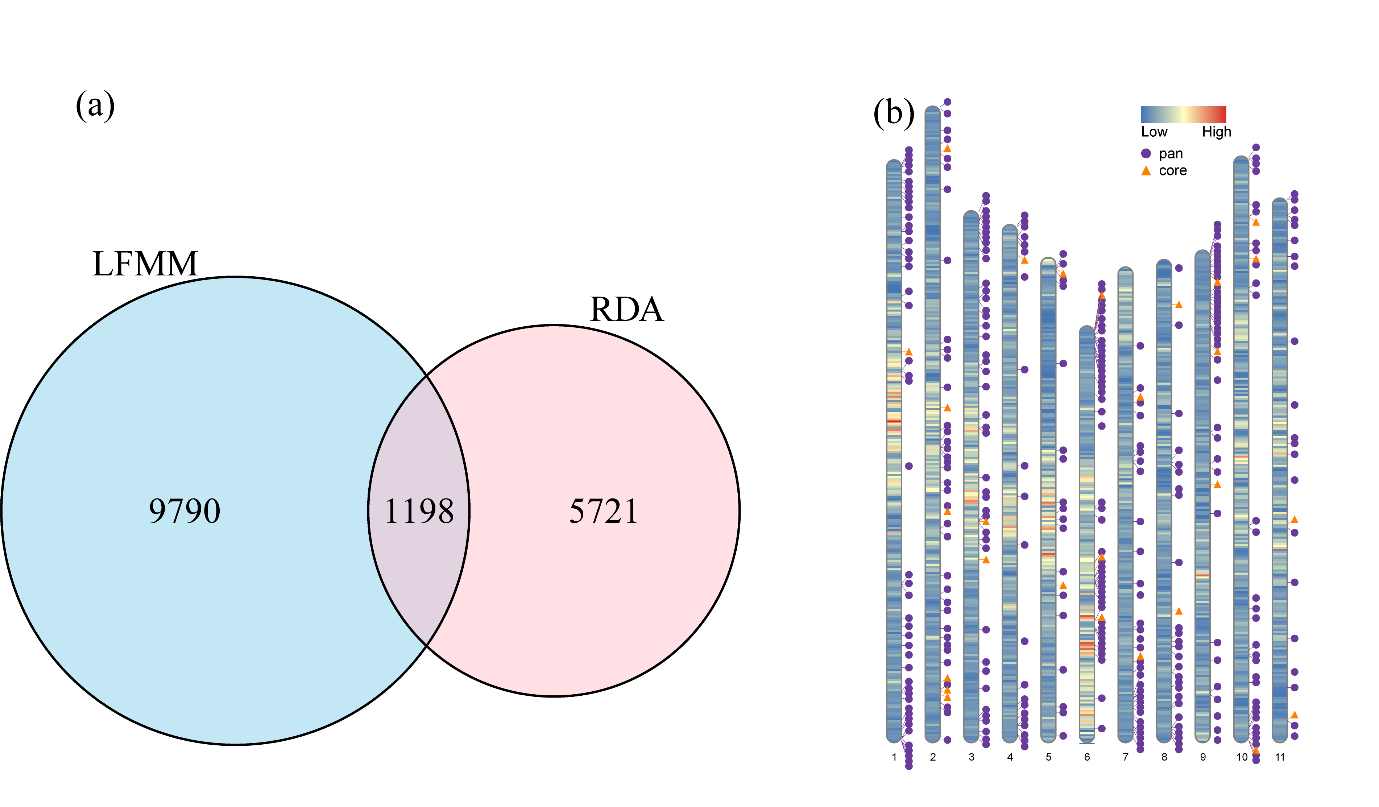

**Supplementary fig. S5** (a) Venn diagram illustrates putatively adaptive SNPs identified within the *Rheum palmatum* complex through latent factor mixed model (LFMM) and redundancy analysis (RDA). (b) Distribution of annotated genes across the whole-genome of *Rheum palmatum*, with adaptive SNP annotation genes corresponding to pan-adaptation loci (purple circles) and core adaptation loci (orange triangles).

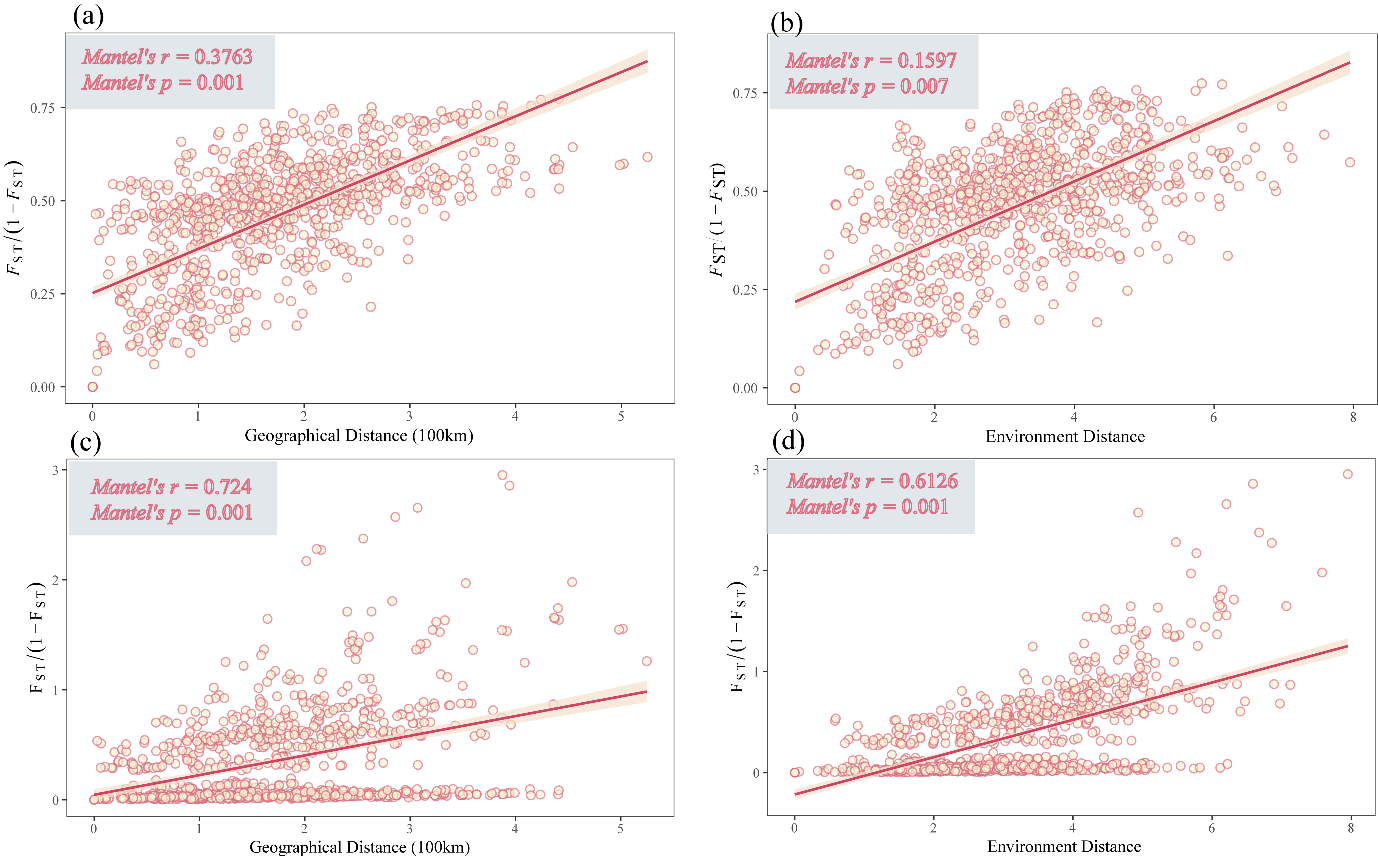

**Supplementary fig. S6** Patterns of genetic differentiation within the *Rheum palmatum* complex. Panels (a) and (c) represent the isolation-by-distance analysis (two-sided Mantel test) for (a) pan adaptation loci and (c) core adaptation. Shadows represent the 95% confidence interval of linear regression. Panels (b) and (d) represent the isolation-by-environment analysis (two-sided partial Mantel test) for (b) pan-adaptation loci and (d) core adaptation loci, with shadows indicating the 95% confidence interval.
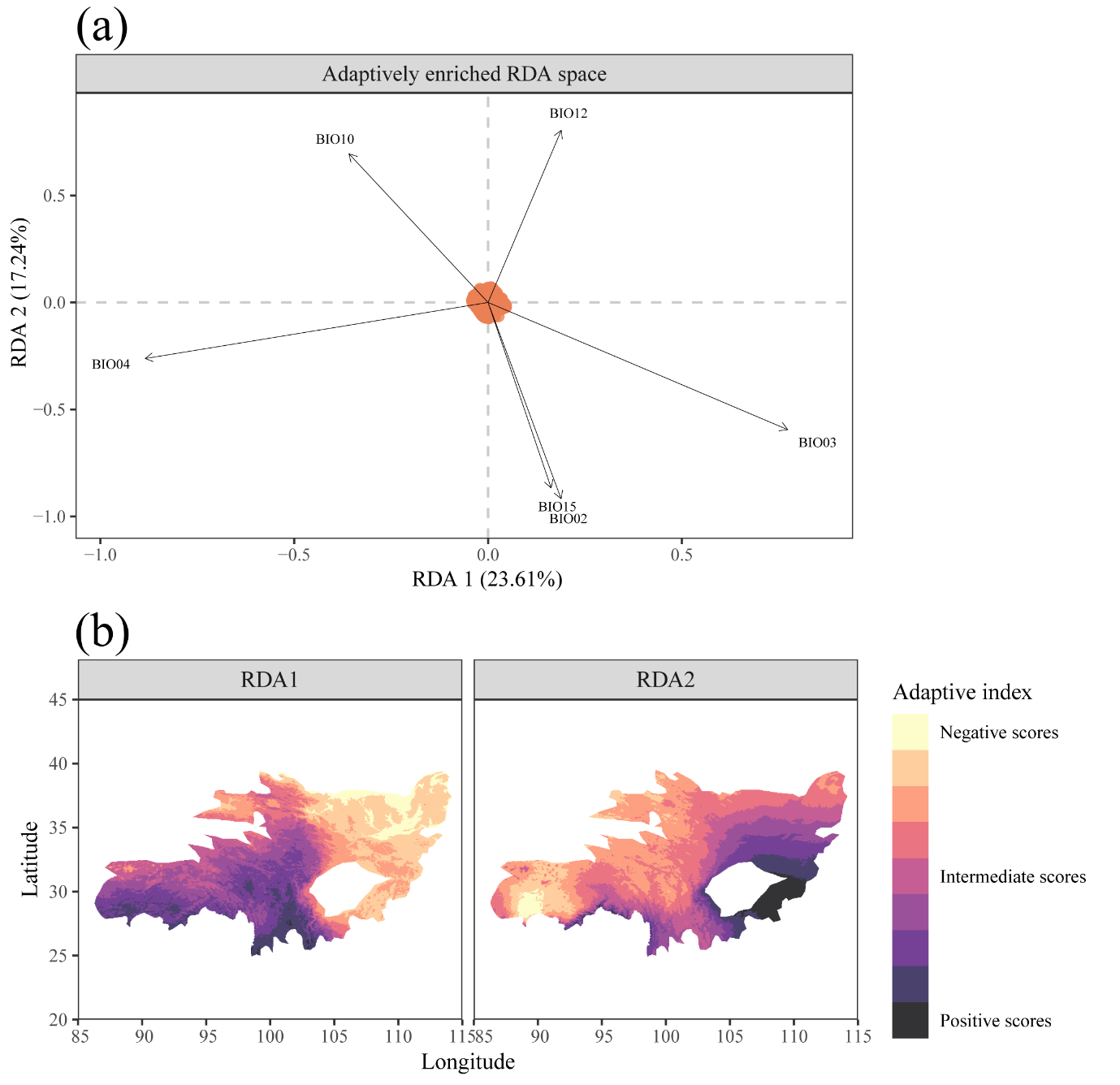

**Supplementary fig. S7** The adaptive landscape of the *Rheum palmatum* complex. (a) The adaptively enriched genetic space showing association between pan-adaptation loci and the six key climatic drivers of adaptation. (b) Spatial projection of adaptive genetic turnover across the potential range of *Rheum palmatum* complex.

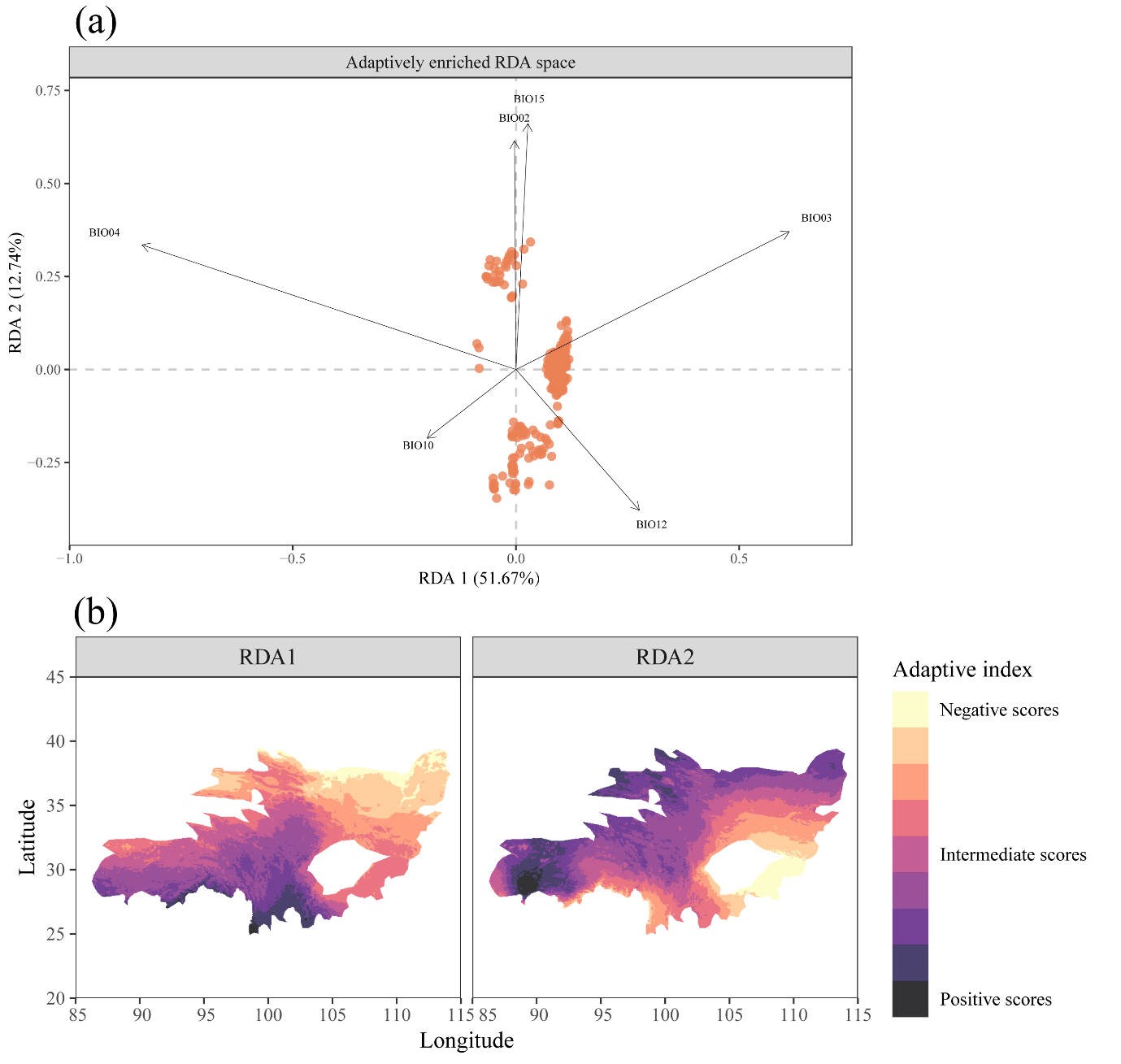

**Supplementary fig. S8** The adaptive landscape of the *Rheum palmatum* complex. (a) The adaptively enriched genetic space showing association between core adaptation loci and the six key climatic drivers of adaptation. (b) Spatial projection of adaptive genetic turnover across the potential range of *Rheum palmatum* complex.

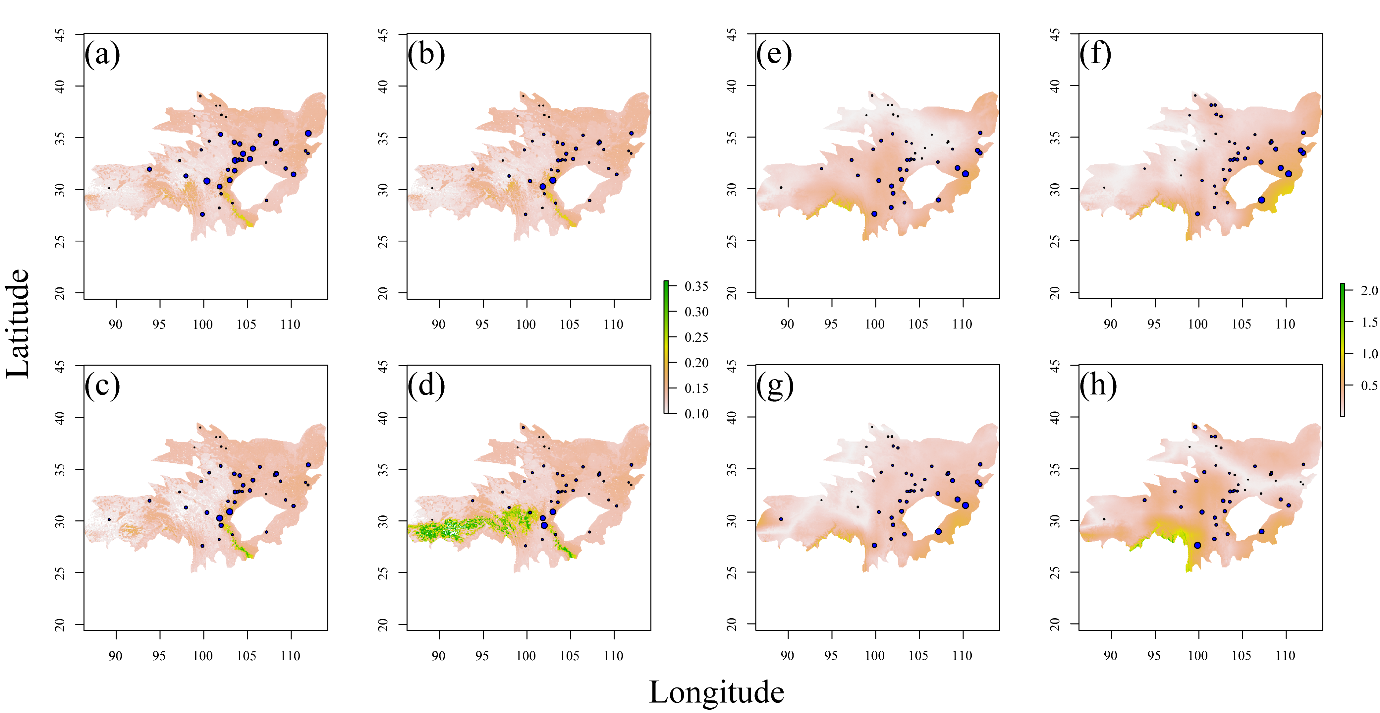

**Supplementary fig. S9** Predicted genomic offsets across the range of *Rheum palmatum* complex in response to future climates, derived from core adaptation loci using gradient forest (panels a-d) and redundancy analysis (panels e-h). Panels (a) and (e) represent the scenarios corresponding to the shared socioeconomic pathways SSP126 in 2050, while (b) and (f) depict SSP126 in 2090. Similarly, (c) and (g) illustrate SSP 585 in 2050, and (d) and (h) show SSP585 in 2090. The size of the blue circles in each panel corresponds to the magnitude of the genomic offset under the respective climate scenarios.

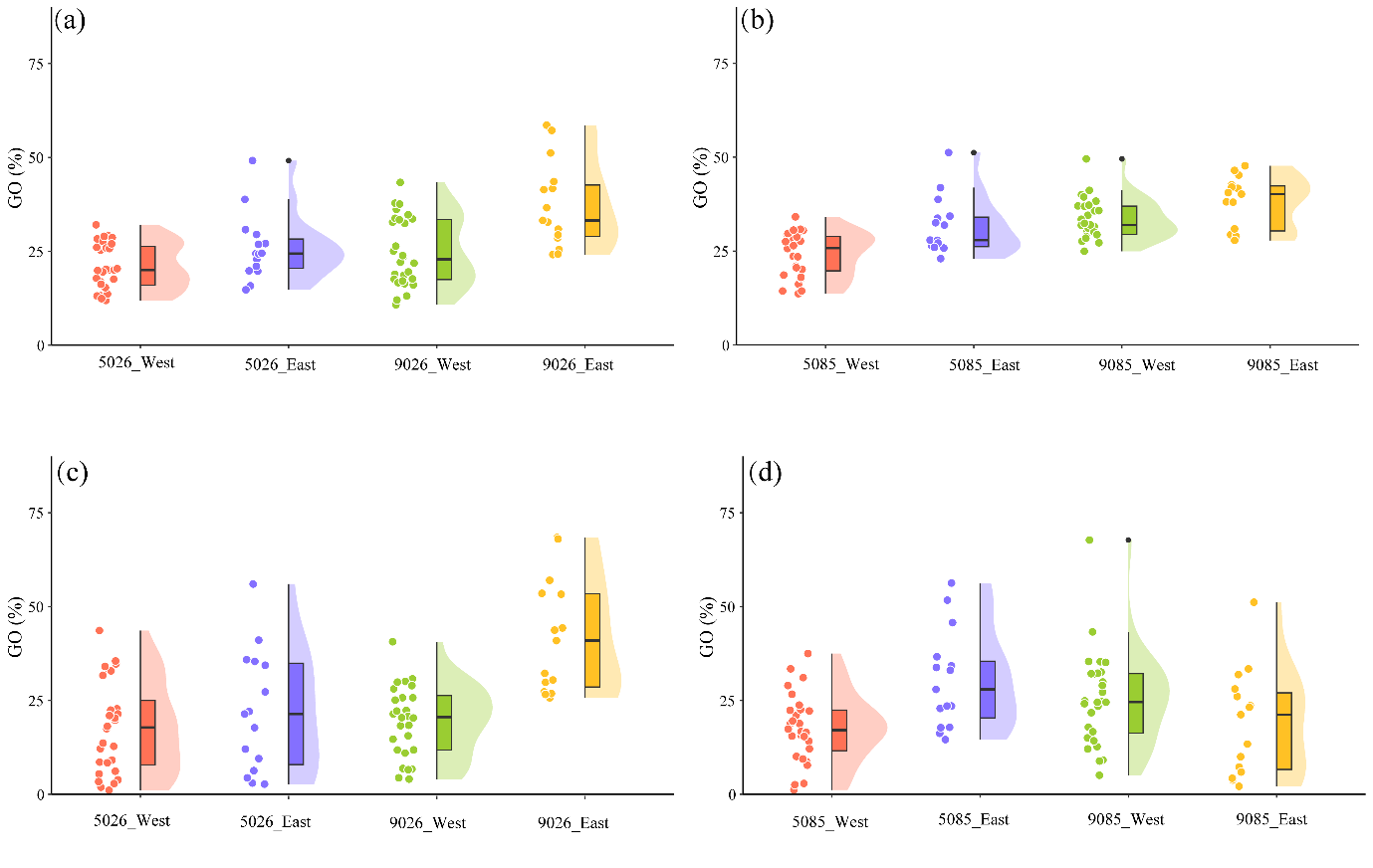

**Supplementary fig. S10** Predicted genomic offsets of the western and eastern lineage populations derived from species-level estimations, utilizing both pan-adaption loci (panels a and b) and core adaptation loci (panels c and d) via redundancy analysis. On the x-axis, the numerals 5026 and 9026 denote the scenarios corresponding to the shared socioeconomic pathways SSP126 in 2050 and 2090, respectively, while 5085 and 9085 represent the SSP585 scenarios in 2050 and 2090, respectively.

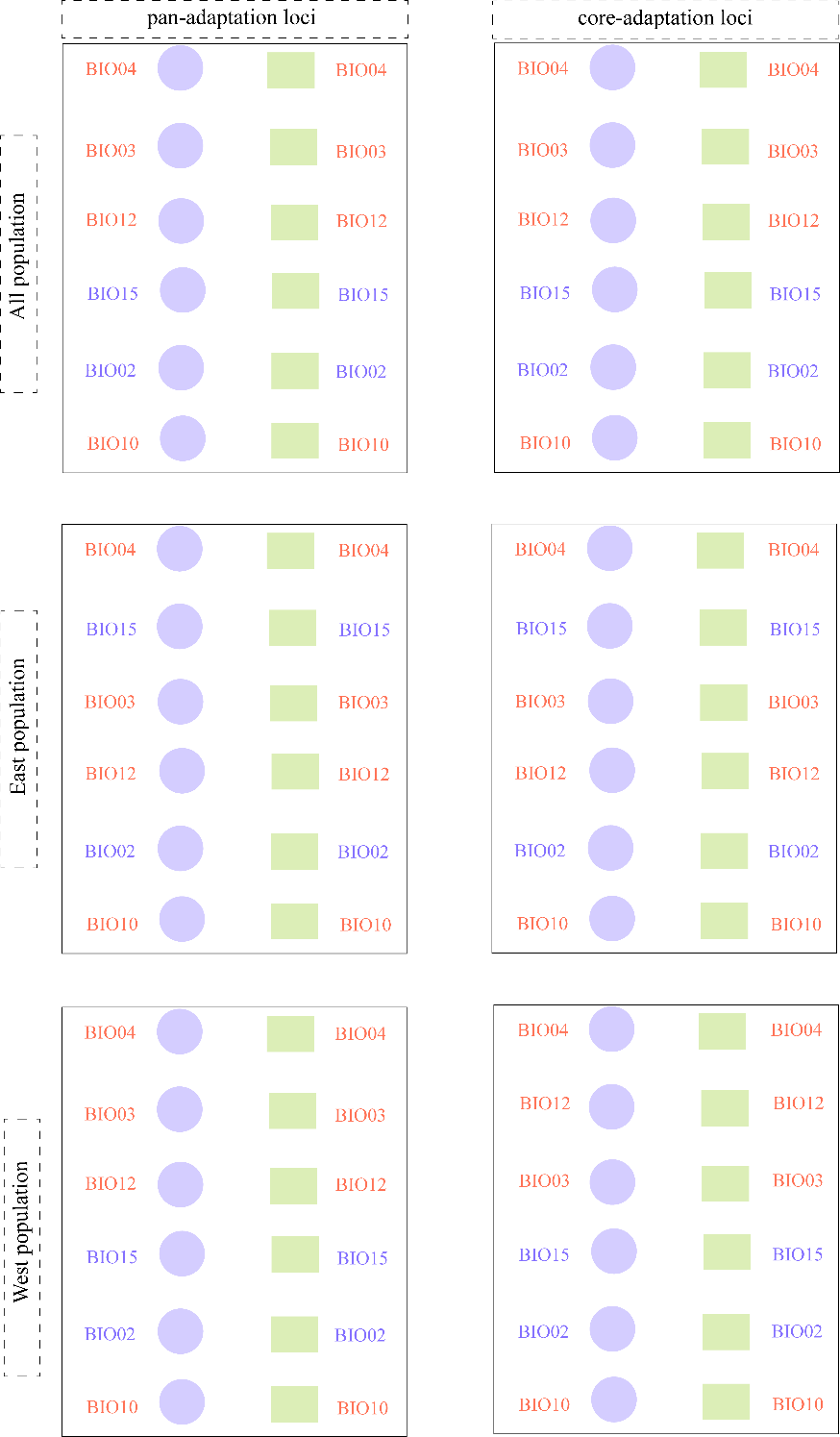

**Supplementary fig. S11** Consistency in environmental accuracy (depicted by light purple circles) and weighted importance rankings (represented by light green rectangles) as outputted by trained gradient forest models utilizing pan-adaptation loci and core adaptation loci at species and lineage levels.

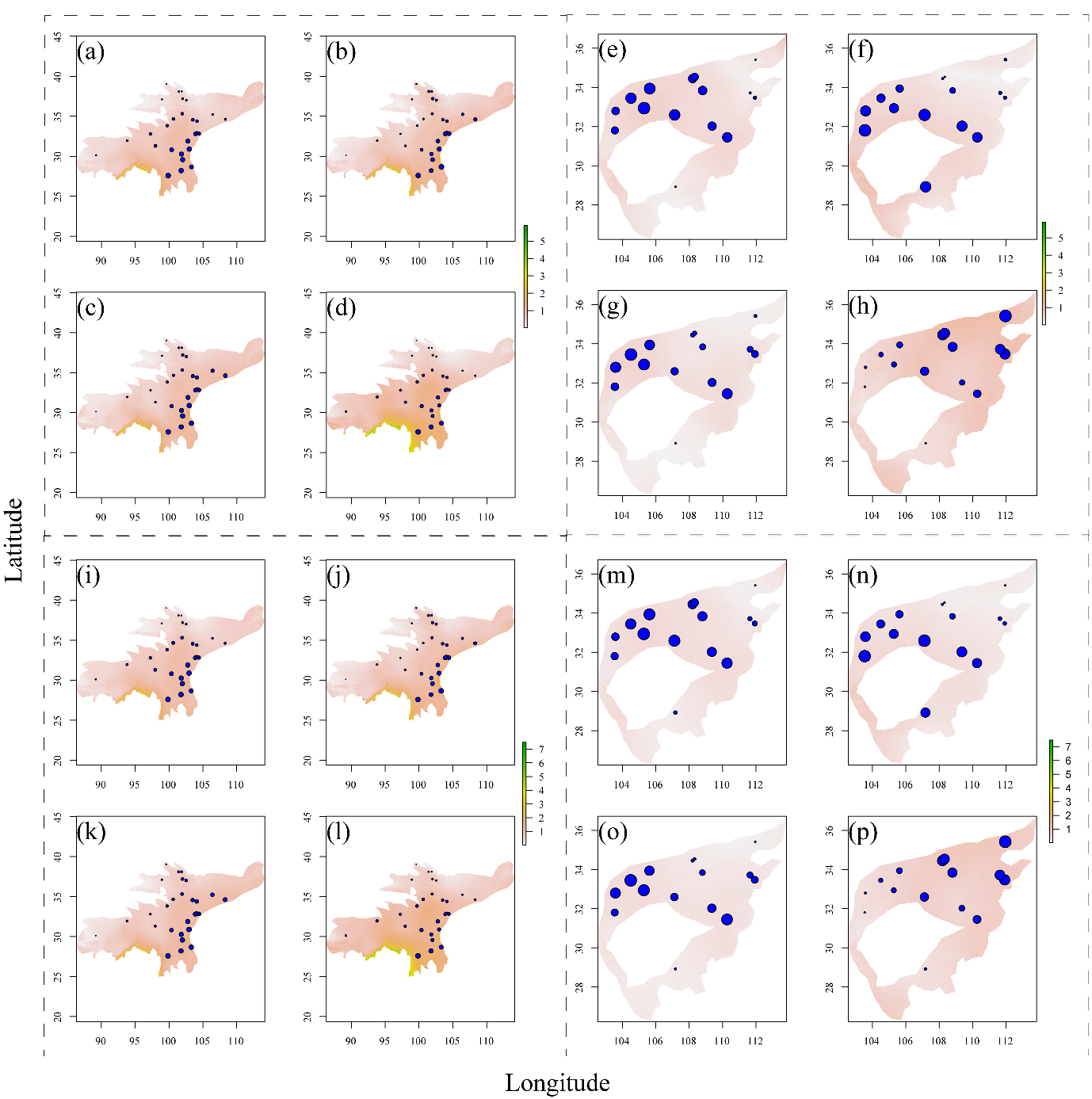

**Supplementary fig. S12** Predicted genomic offsets at the lineage level based on pan-adaptation loci (panels a-h) and core adaptation loci (panels i-p) using redundancy analysis. Panels (a) and (i) depict the offsets for western lineage populations under the scenarios corresponding to the shared socioeconomic pathways SSP126 in 2050, while (b) and (j) represent the same for 2090. Panels (c) and (k) exhibit offsets for SSP585 in 2050, and (d) and (l) for 2090. For eastern lineage populations, panels (e) and (m) display offsets under SSP126 in 2050, (f) and (n) for 2090, (g) and (o) for SSP585 in 2050, and finally, (h) and (p) for 2090.

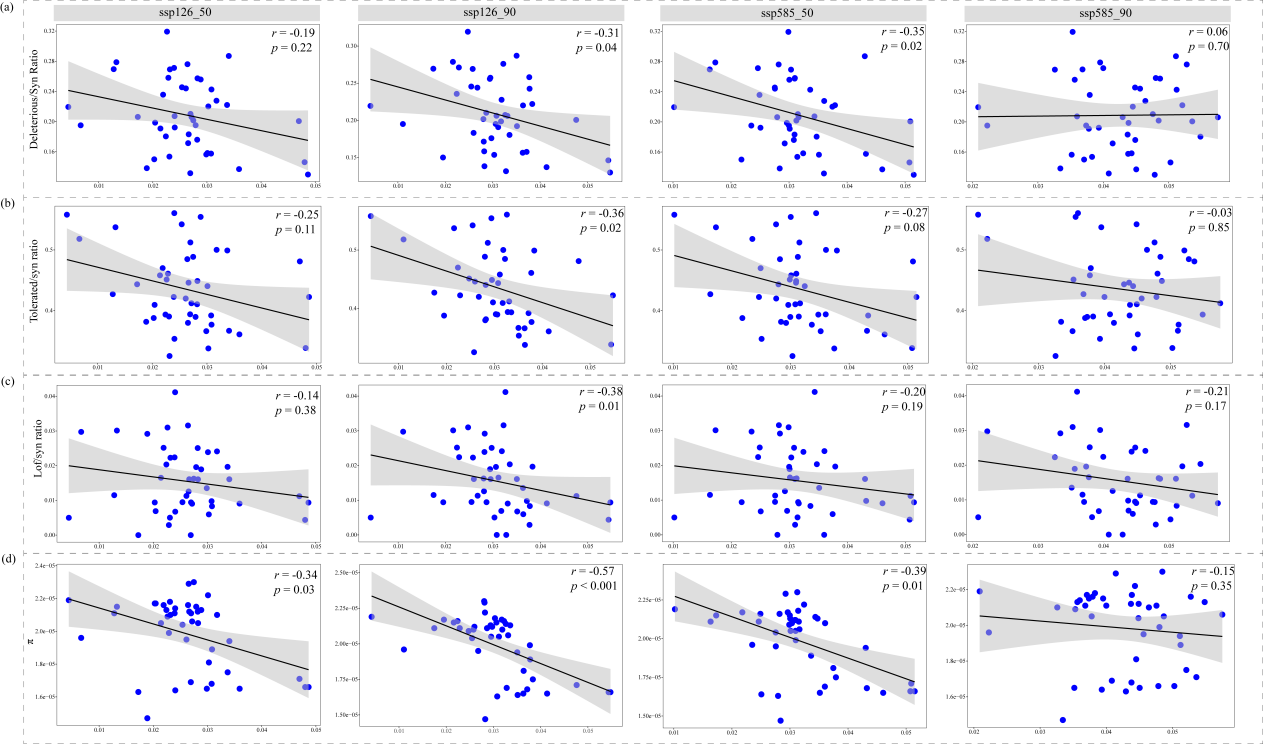

**Supplementary fig. S13** Relationships between various proxies of genetic load (y-axis) and genomic offset (x-axis) via gradient forest based on pan-adaptation loci under SSP126 socioeconomic pathway scenarios for 2050 (a) and 2090 (b), and SSP585 scenarios for 2050 (c) and 2090 (d). (a) Relationship between genomic offset and the ratio of derived deleterious variants to derived synonymous variants. (b) Relationship between genomic offset and the ratio of derived tolerant variants to derived synonymous variants. (c) Relationship between genomic offset and the ratio of loss-of-function variants to derived synonymous variants. (d) Relationship between genomic offset and nucleotide diversity.

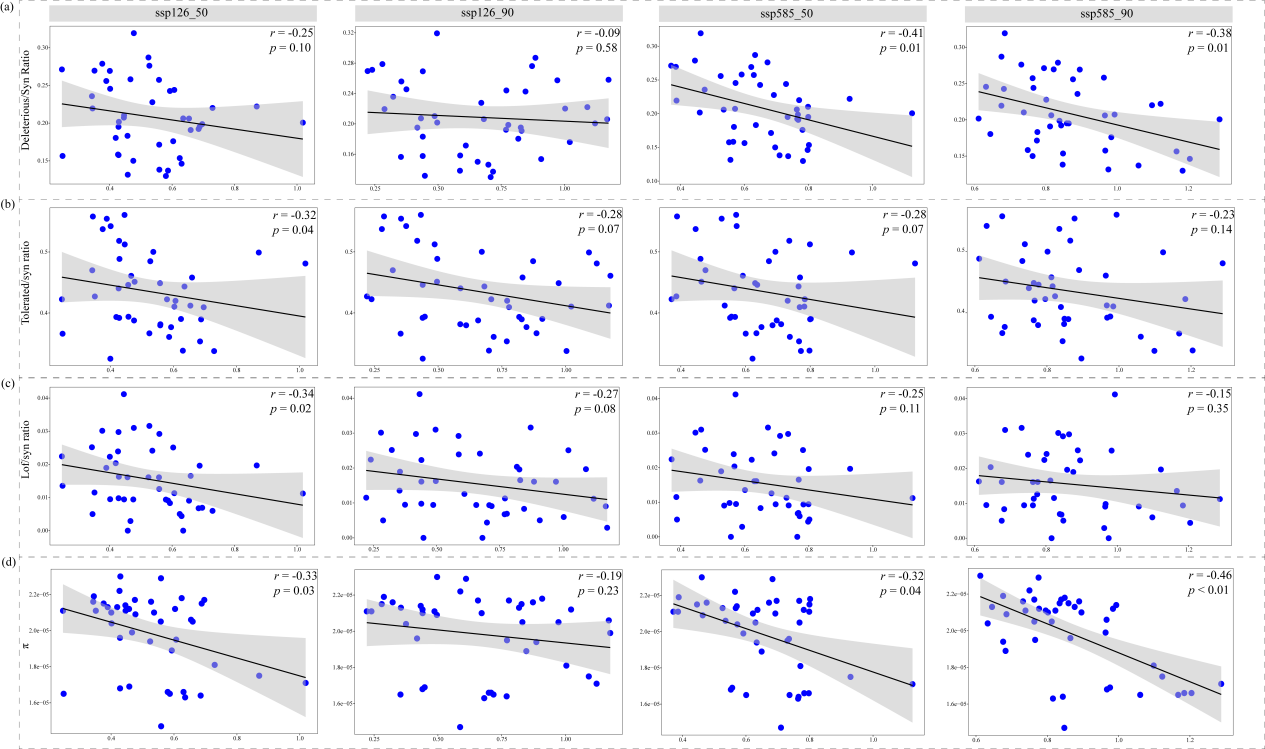

**Supplementary fig. S14** Relationships between various proxies of genetic load (y-axis) and genomic offset (x-axis) via redundancy analysis based on pan-adaptation loci under SSP126 socioeconomic pathway scenarios for 2050 (a) and 2090 (b), and SSP585 scenarios for 2050 (c) and 2090 (d). (a) Relationship between genomic offset and the ratio of derived deleterious variants to derived synonymous variants. (b) Relationship between genomic offset and the ratio of derived tolerant variants to derived synonymous variants. (c) Relationship between genomic offset and the ratio of loss-of-function variants to derived synonymous variants. (d) Relationship between genomic offset and nucleotide diversity.

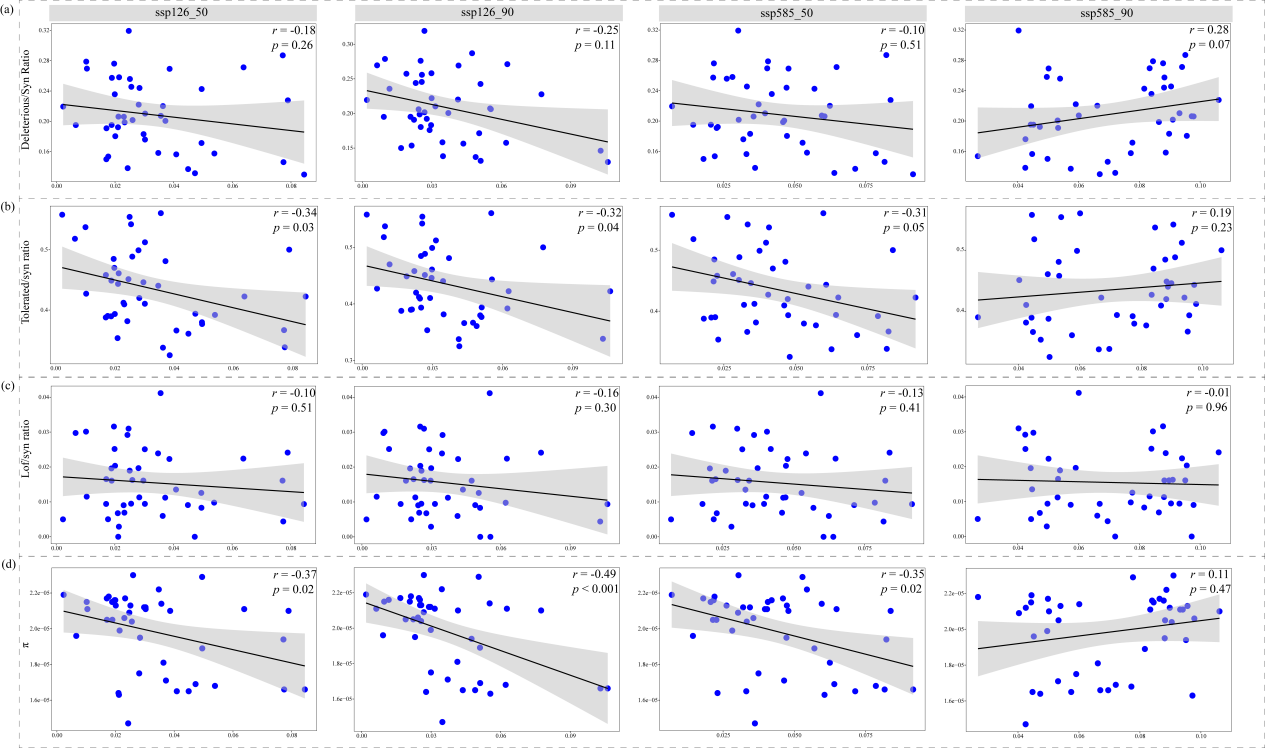

**Supplementary fig. S15** Relationships between various proxies of genetic load (y-axis) and genomic offset (x-axis) via gradient forest based on core adaptation loci under SSP126 socioeconomic pathway scenarios for 2050 (a) and 2090 (b), and SSP585 scenarios for 2050 (c) and 2090 (d). (a) Relationship between genomic offset and the ratio of derived deleterious variants to derived synonymous variants. (b) Relationship between genomic offset and the ratio of derived tolerant variants to derived synonymous variants. (c) Relationship between genomic offset and the ratio of loss-of-function variants to derived synonymous variants. (d) Relationship between genomic offset and nucleotide diversity.

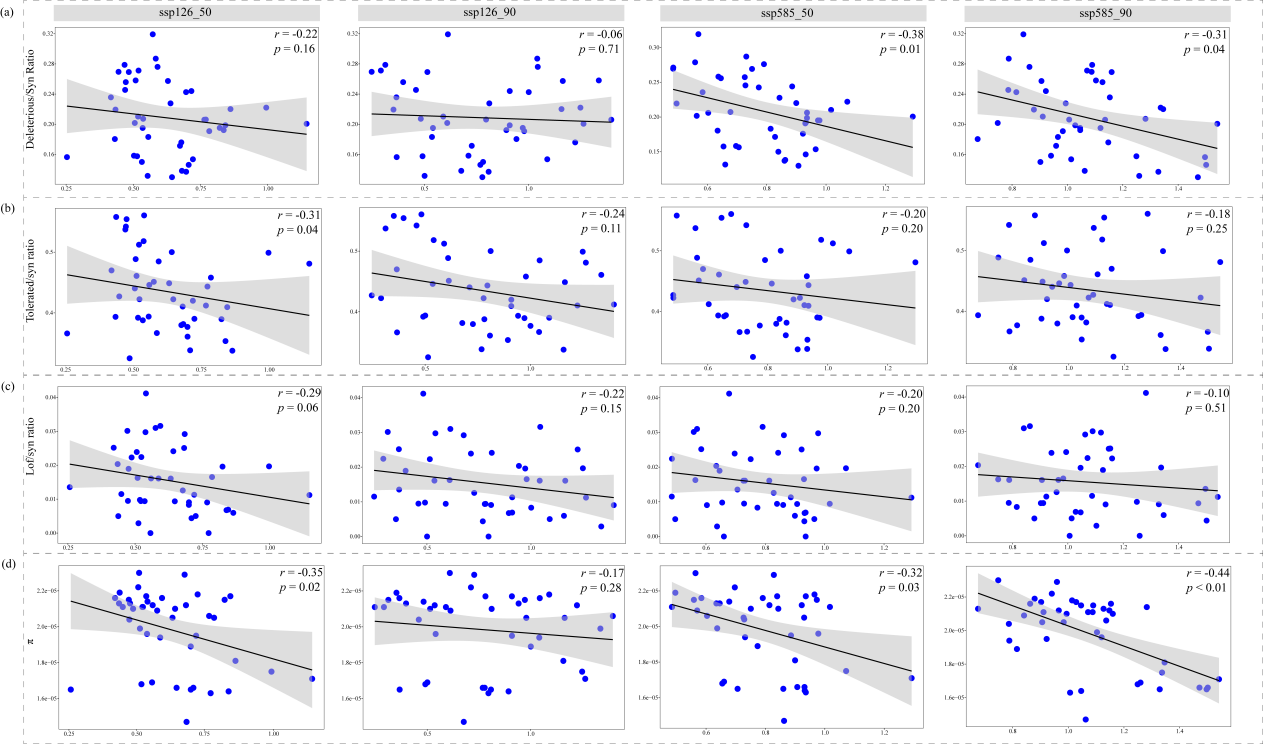

**Supplementary fig. S16** Relationships between various proxies of genetic load (y-axis) and genomic offset (x-axis) via redundancy analysis based on core adaptation loci under SSP126 socioeconomic pathway scenarios for 2050 (a) and 2090 (b), and SSP585 scenarios for 2050 (c) and 2090 (d). (a) Relationship between genomic offset and the ratio of derived deleterious variants to derived synonymous variants. (b) Relationship between genomic offset and the ratio of derived tolerant variants to derived synonymous variants. (c) Relationship between genomic offset and the ratio of loss-of-function variants to derived synonymous variants. (d) Relationship between genomic offset and nucleotide diversity.

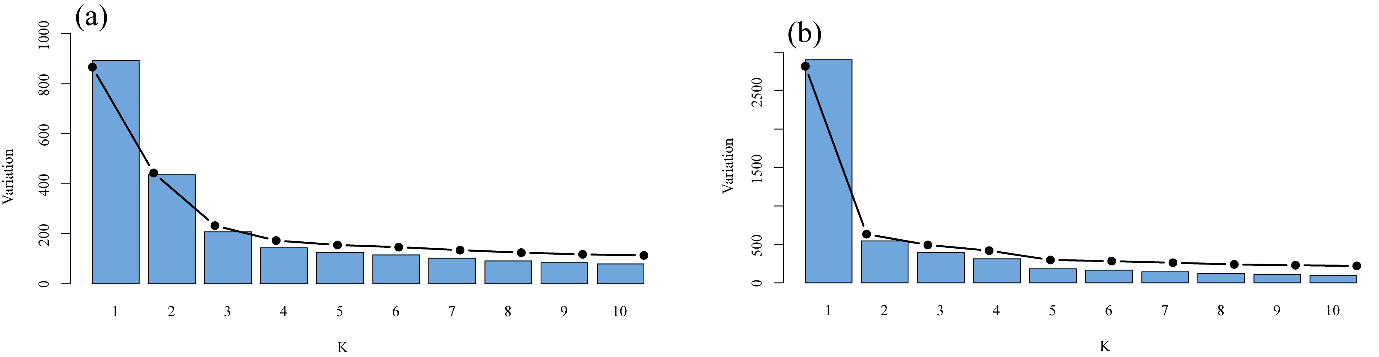

**Supplementary fig. S17** Intra-cluster variation (black solid line) and its reduction (blue bars) with increasing number of clusters, as estimated from continuous genomic variation predicted by gradient forest based on pan-adaptation loci (a) and core adaptation loci (b) across the range of *Rheum palmatum* complex.

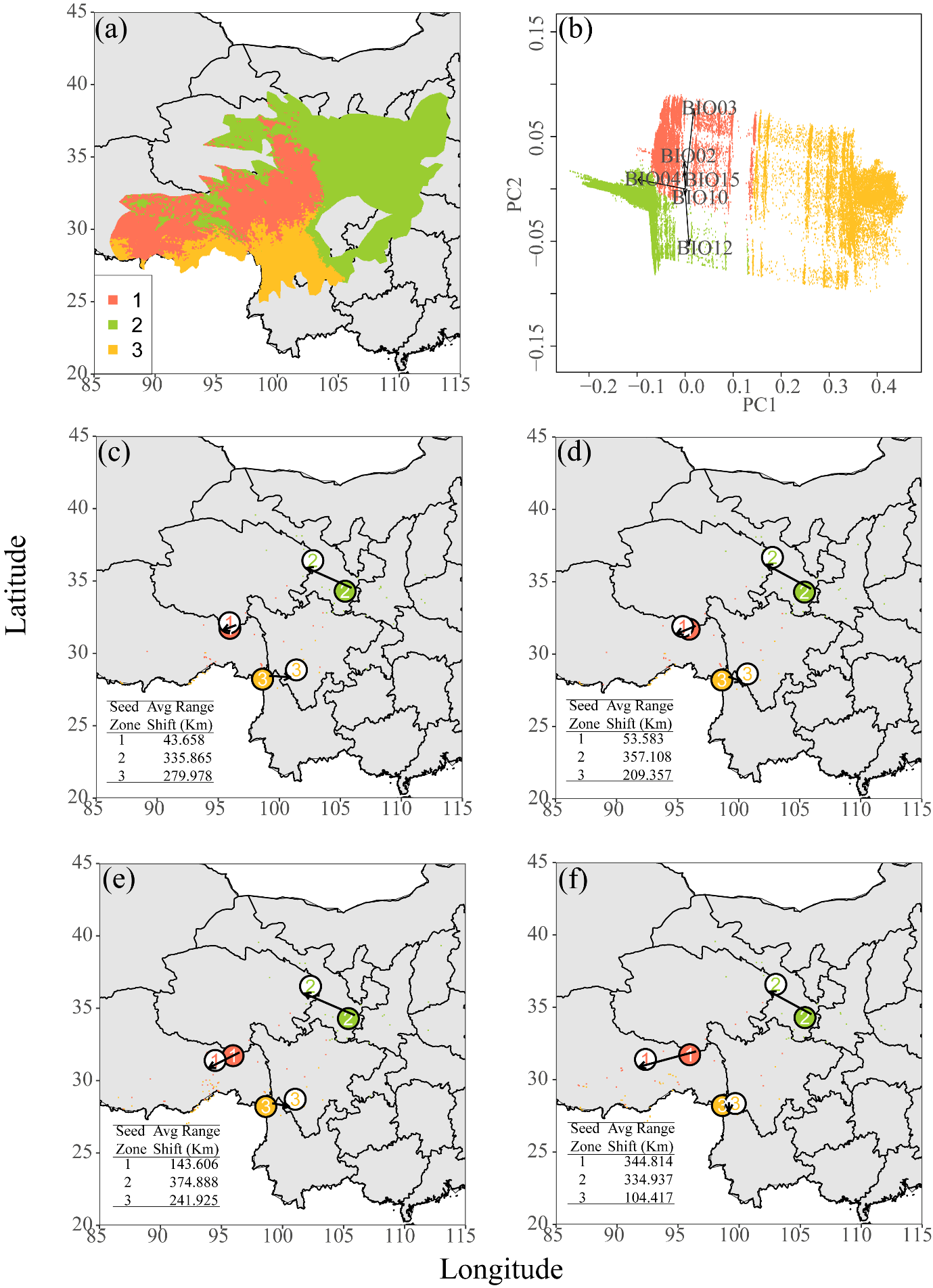

**Supplementary fig. S18** Predicted seed zones based on core adaptation loci and their range shifts to minimize genomic offset under climate change. Panel (a) represents predicted three seed zones for the *Rheum palmatum* complex using gradient forest (GF). The biplot in subfigure illustrates the GF-predicted genomic variation used for seed zone delineation. Panels (b-e) represent the predicted range adjustments for each seed zone to minimize genomic disparities derived from pan-adaptation loci under various climate scenarios. Solid-colored numerals denote the current centroids of the seed zones, whereas numerals on a white background indicate the anticipated centroid shifts in alignment with future climate projections. Panels (b) and (c) correspond to the SSP126 socioeconomic pathway scenarios in 2050 and 2090, respectively. Panels (d) and (e) reflect the SSP585 scenarios in 2050 and 2090, respectively. The colored pixels represent the predicted relocation points for each seed zone, where genomic disparities are minimized based on future climate forecasts (red = Seed Zone 1, yellow = Seed Zone 2, green = Seed Zone 3). The inset table clarifies the distance in kilometers between the present and predicted centroids for each seed zone.
